# Hornworts reveal a spatial model for pyrenoid-based CO_2_-concentrating mechanisms in land plants

**DOI:** 10.1101/2024.06.26.600872

**Authors:** Tanner A. Robison, Zhen Guo Oh, Declan Lafferty, Xia Xu, Juan Carlos A. Villarreal, Laura H. Gunn, Fay-Wei Li

**Affiliations:** Boyce Thompson Institute, Ithaca, NY 14853, USA; Plant Biology Section, Cornell University, Ithaca, NY 14853, USA; Department of Biology, Laval University, Quebec City, QC, Canada

## Abstract

Pyrenoid-based CO_2_-concentrating mechanisms (pCCMs) turbocharge photosynthesis by saturating CO_2_ around Rubisco. Hornworts are the only land plants with a pCCM. Owing to their closer relationship to crops, hornworts could offer greater translational potential compared to the green alga Chlamydomonas, the traditional model for studying pCCM. Here we report the first thorough investigation of a hornwort pCCM using the emerging model *Anthoceros agrestis*. The pyrenoids in *A. agrestis* exhibit liquid-like properties similar to Chlamydomonas, but differ by lacking starch sheaths and being enclosed by multiple thylakoids. We found that the core pCCM components in Chlamydomonas, including BST, LCIB, and CAH3, are conserved in *A. agrestis* and likely have similar functions based on their subcellular localizations. Therefore, the underlying chassis for concentrating CO_2_ might be shared between hornworts and Chlamydomonas, and ancestral to land plants. Our study presents the first spatial model for pCCM in a land plant, paving the way for future biochemical and genetic investigations.

## MAIN

Ribulose-1,5-bisphosphate carboxylase/oxygenase (Rubisco), the carbon-fixing enzyme, is the gatekeeper for virtually all biologically available carbon. Despite its central importance in global primary productivity, Rubisco is considered to have two major limitations: a low rate of activity and poor specificity for carbon dioxide (CO_2_) ^1^. Photosynthetic organisms have largely overcome the first of these limitations by simply making more of the enzyme, so much so that Rubisco is considered to be the most abundant enzyme in the biosphere ^2^. Rubisco’s poor specificity means that it can also react with oxygen (O_2_), resulting in photorespiration which costs energy and leads to the net loss of fixed CO_2_ ^3,4^.

To reduce photorespiration, some plants have evolved systems to concentrate inorganic carbon (Ci) around Rubisco (called CO_2_-concentrating mechanisms; CCM), either through biophysical or biochemical means ^5^. Biochemical CCMs (C_4_ or CAM photosynthesis) use enzymatic pathways to concentrate carbon, either in vacuoles or in bundle sheath cells.

Biophysical CCMs, seen in cyanobacteria, algae, and hornworts, concentrate Ci in subcellular compartments (pyrenoids or carboxysomes) where Rubisco is highly abundant ^6^.

The model organism for studying pyrenoid-based CCM (pCCM) is the green alga *Chlamydomonas reinhardtii*. Chlamydomonas pyrenoids are a liquid-like proteinaceous compartment whose phase separation is mediated by the interaction of Rubisco and the linker protein EPYC1 ^7–9^. In the algal pCCM, a series of Ci channels, such as LCI1, LCIA, and bestrophin channels (BST), facilitate Ci transport from the extracellular space into specialized thylakoid tubules ^10^. These tubules traverse pyrenoids and contain a specific carbonic anhydrase (CA), CAH3. This CA catalyzes the conversion of bicarbonate (HCO_3_^-^) into CO_2_, which can freely diffuse out of the thylakoid tubules and into the pyrenoid. A starch sheath surrounds pyrenoids and might function as a CO_2_ diffusion barrier to enhance the efficiency of the CCM ^11,12^. Another CA, LCIB, localizes around the gaps of the starch sheath to recapture leaked CO_2_ by converting it back to HCO_3_^-^. Recent modeling work suggests that a minimally functional pCCM requires the joint operation of BST, LCIB, and CAH3 (or the equivalent thereof), in addition to a pyrenoid enclosed by diffusion barriers ^13^.

Installing a pCCM into crop plants may boost CO_2_ fixation by as much as 60% ^4,14,15^.

However, transplanting an algal CCM into land plants is complicated by the fact that around one billion years of evolution separate these lineages ^16^, over which time significant differences have accumulated in chloroplast structure and protein sequences. On the other hand, hornworts, a group of bryophytes (Fig. 1A,B), are the only known land plants with a pyrenoid or biophysical CCM of any kind ^17–19^. Hornwort pyrenoids are functionally analogous to algal pyrenoids, acting as a locus of Rubisco accumulation, and the focal point of the pCCM ^17,18,20^. Characterizing the land plant pCCM may provide significant translational advantages. However, virtually nothing is known about the functional components enabling a pCCM in hornworts.

**Figure 1.**
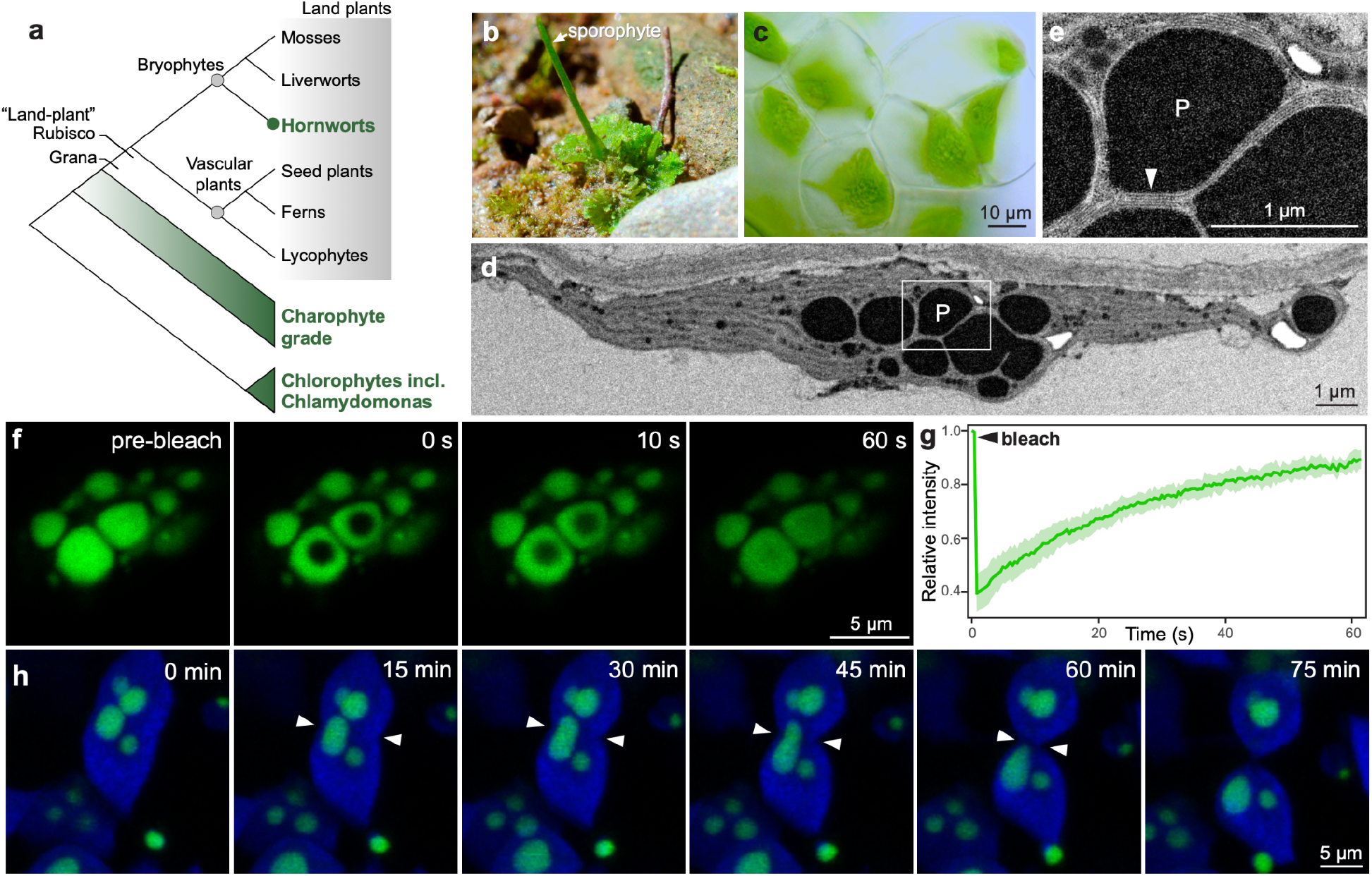
Morphology and physical property of pyrenoids in *Anthoceros agrestis*. (a) Hornworts are the only land plants that have a pCCM. Compared to Chlamydomonas, hornworts and crop plants share more common features in their chloroplast and Rubisco structures. (b) The model hornwort *Anthoceros agrestis* in native habitat. (c) *A. agrestis*, like many other hornwort species, has a single chloroplast per cell. (d) TEM image of an *A. agrestis* chloroplast with multiple pyrenoids, P. (e) A close-up view of a pyrenoid from (D). The arrowhead points to a stack of thylakoid membranes encasing the pyrenoid. (f) FRAP of hornwort pyrenoids labeled with RCA::mVenus. Photobleaching was targeted at the center of two selected pyrenoids. (scale bar, 5 µm.) (g) FRAP recovery curve. Error bars represent standard error of the mean (SEM) and are shown as the green shaded area (n=6). (h) Time-lapse images during cell division, showing signals of chlorophyll autofluorescence (blue) and RCA::mVenus fluorescence (green) over 75 mins. The cell division plane is demarcated with white arrowheads. (scale bar, 5 µm.)

In this study, we characterized the morphology and physical properties of hornwort pyrenoids, as well as proteins involved in the pCCM, using the emerging model hornwort *Anthoceros agrestis* ^21–23^. Diffusion properties of fluorescently tagged proteins and live-cell imaging indicate that *A. agrestis* pyrenoids are liquid-like, similar to the Chlamydomonas pyrenoid. Putative CCM components for protein subcellular localization studies were selected by leveraging both Rubisco co-immunoprecipitation (co-IP) and 41 genomes spanning plant and algal diversity. We found that hornworts possess orthologs of several core pCCM components from Chlamydomonas, and provide evidence for shared functional roles. We thus infer that the chassis for pCCM was present in the last common ancestor of land plants. Comparative genomics and co-IP did not reveal any clear EPYC1 homolog or analog, implying that hornworts might have adopted a different strategy for pyrenoid formation. Based on our findings, we propose the first spatial model for a land plant pCCM, which is consistent with reaction-diffusion modeling for a functional pCCM, and set the stage for future biochemical and genetic investigation.

## RESULTS

### *A. agrestis* contains multiple pyrenoids that are not enclosed by starch sheaths

We first characterized the morphology of *A. agrestis* pyrenoids using transmission electron microscopy (TEM). Compared to other hornwort species ^17,20^, *A. agrestis* pyrenoids are slightly larger in size, with a diameter ranging from 500 nm to 5 μm (Fig. 1c-e; Supplementary Fig. 1). *A. agrestis* (and hornworts in general) have multiple pyrenoids per chloroplast, though the number of which appears to be highly variable (Fig. 1c,d; Supplementary Fig. 1). This pattern differs from the singular pyrenoid found in most algal species including Chlamydomonas ^24^. Further, we found a clear lack of any kind of starch sheath surrounding the pyrenoids of *A. agrestis* (Fig. 1d,e), which in Chlamydomonas is believed to act as a critical diffusion barrier to prevent CO_2_ leakage ^11,12^. While no starch sheath is present, *A. agrestis* has many layers of stacked thylakoids wrapping around the pyrenoids (Fig. 1e; white arrowhead). Such stacked thylakoids might be sufficient to prevent CO_2_ leakage, as suggested by recent modeling of the Chlamydomonas pCCM ^13^.

### *A. agrestis* pyrenoids are liquid-like

While we have previously demonstrated that the pyrenoids of *A. agrestis* are the sites of Rubisco accumulation ^23^, the physical properties of these organelles have not been investigated. Chlamydomonas pyrenoids are known to be liquid-like in nature, whereas the pyrenoids of the diatom *Phaeodactylum tricornutum* exhibit much less internal mixing ^25^. To infer the physical properties of *A. agrestis* pyrenoids, we applied the fluorescence recovery after photobleaching (FRAP) technique on our stable transgenic line expressing Rubisco activase (RCA) tagged with mVenus. RCA was previously shown to co-localize with Rubisco in *A. agrestis* ^23^, and can thus be used as a pyrenoid marker. We found that the mVenus signal exhibited a rapid recovery (t_1/2_ = 16.2 s) post photobleaching (Fig 1f,g). The unbleached region equilibrates with the bleached region of the same pyrenoid within 60 s (Fig. 1f,g), suggesting high levels of internal mixing. Furthermore, live cell imaging of dividing chloroplasts in the same RCA::mVenus line demonstrated elongation of a pyrenoid as it is being pulled to one of the daughter cells, again suggesting that *A. agrestis* pyrenoids are liquid-like (Fig. 1h, Movie S1).

### Photosystems I and II are both in close proximity to pyrenoids

Given the tight association between thylakoids and pyrenoids in hornworts (Fig. 1d,e), we next examined how thylakoids are distributed across the chloroplasts of *A. agrestis*, especially whether there was differential localization of Photosystem I (PSI) and Photosystem II (PSII) relative to the pyrenoids. It is thought that in hornworts PSII localizes mainly to the grana, while PSI has a more ubiquitous distribution including in the channel thylakoids ^20,26^. The evidence supporting this claim is, however, limited. To visualize PSI and PSII distribution, we transiently expressed fluorescently tagged Photosystem I reaction center subunit VI (PSAH) and Oxygen-evolving enhancer protein 2 (PSBP), respectively, in *A. agrestis* using a biolistics transformation approach ^23^.

In agreement with TEM images (Fig. 1d,e), confocal imaging of tagged PSAH and PSBP showed a tight association between thylakoids and pyrenoids (Fig. 2a,b; Supplementary Fig. 2). Pyrenoids are almost entirely enveloped by thylakoids (Fig. 2a,b), and both PSI and PSII localize in the membranes directly adjacent to the pyrenoids (Fig. 2a,b, d,e). Consistent with previous reports ^20^, PSI is uniformly distributed across the thylakoid network in hornworts, so much so that the signal is only absent in pyrenoid bodies (Fig. 2a; Supplementary Fig. 2). Localization of PSII appears to be less uniform than PSI, with gaps as well as smaller regions of much higher intensity, likely grana, scattered throughout the chloroplast (Fig. 2b; Supplementary Fig. 2).

**Figure 2.**
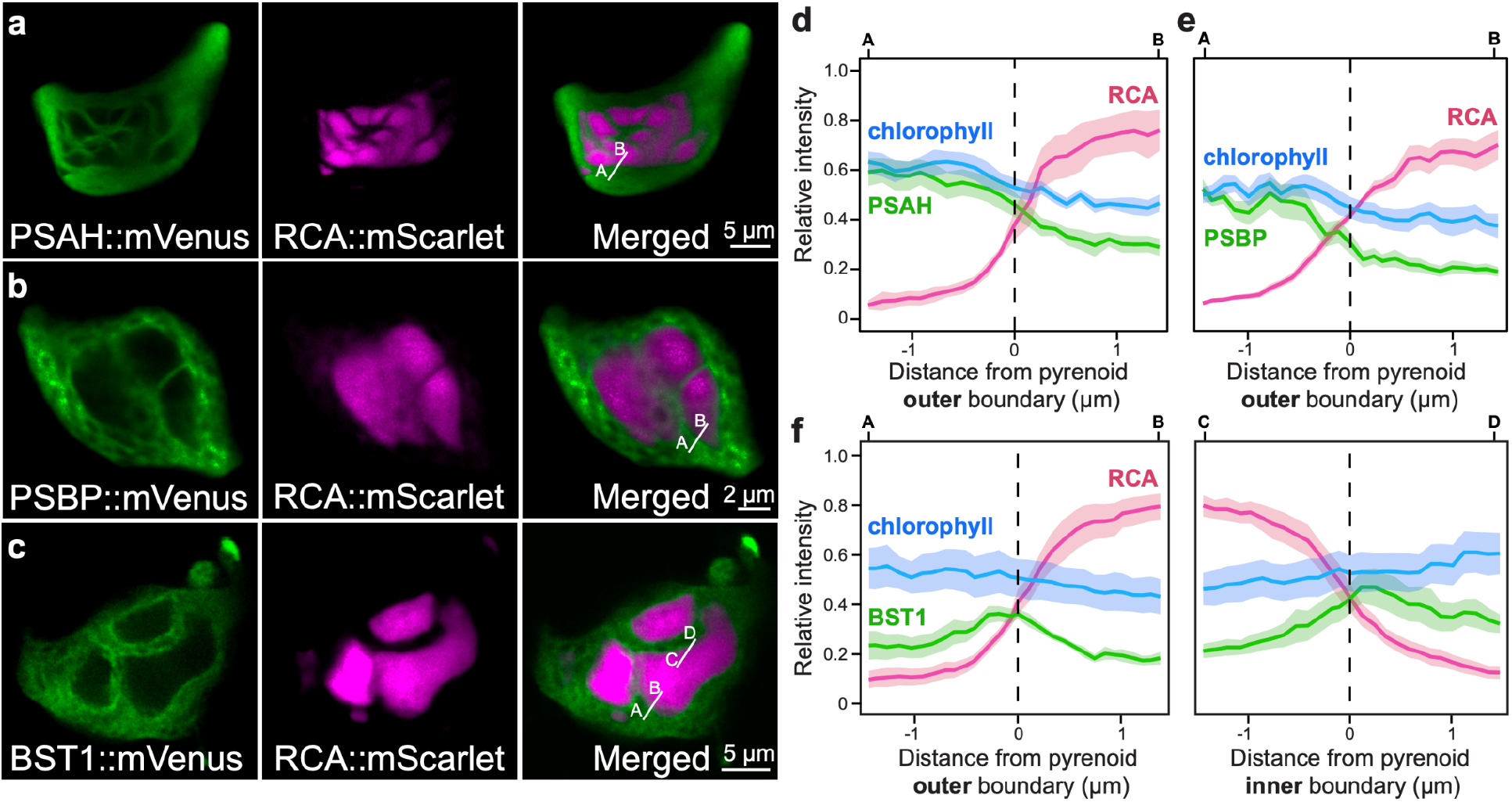
Distribution of photosystems and BST channels on thylakoids of *A. agrestis*. (a) Photosystem I (marked by PSAH::mVenus) and (b) photosystem II (marked by PSBP::mVenus) are distributed throughout the thylakoid network. (scale bar, 2 µm.) (c) BST1::mVenus has elevated fluorescent signals on the thylakoids immediately adjacent to pyrenoids. (scale bar, 5 µm). The fluorescent intensities of (d) PSAH::mVenus and (e) PSAH::mVenus decline along the transects going into pyrenoids (white lines in a and b). (f) Quantification of elevated BST1::mVenus fluorescent signals around pyrenoids (white lines in c). RCA::mScarlet was used as a pyrenoid marker. Error bars represent SEMs and are shown as shaded areas in each curve (n = 10, 11, and 7 for BST, PSII, and PSI, respectively).

### Bestrophin channels localize to thylakoids adjacent to pyrenoids

A biophysical CCM in eukaryotes requires the presence of thylakoid membrane channels to allow HCO_3_^-^ to enter the thylakoid lumen. In Chlamydomonas, a number of bestrophin channels (BST) fill this role ^27^. There are four Chlamydomonas BST orthologs in the *A. agrestis* genome, but only one, BST1, is expressed abundantly (Supplementary Table 1). Fluorescent protein tagging indicated that BST1 in *A. agrestis* was distributed throughout the thylakoid network, but exhibited elevated fluorescence intensity immediately adjacent to the pyrenoids (Fig. 2c,f; Supplementary Fig. 3). This pattern is in contrast to that of PSAH and PSBP, where the fluorescence intensities continuously decline going into pyrenoids (Fig. 2d,e). The localization of *A. agrestis* BST1 is similar to what was reported in Chlamydomonas ^27^, and implies that BST1 likely has the same function in hornworts, transporting HCO_3_^-^ to thylakoid lumen.

### β-CA1 localizes to the cytoplasm

We next examined the localization of several carbonic anhydrases (CA) in *A. agrestis*. There are three known CA families in land plants (α, β, and γ), which catalyze the reversible hydration of CO_2_ to HCO_3_^-28^. While hornworts possess multiple CAs, we focused on three—β-CA1, CAH3, and LCIB—that have high potential to influence pCCM. β-CA1 is the second most abundant protein in plants after Rubisco and is typically located in the chloroplast stroma ^29^.

However, the presence of a CA in the stroma, and specifically the pyrenoid matrix, in hornwort chloroplasts would likely convert CO_2_ into HCO_3_^-^, thus decreasing the availability of CO_2_ for Rubisco and short circuiting the pCCM. The β-CA1 in *A. agrestis*, unlike the orthologs in C_3_ plants, lacks a predicted chloroplast transit peptide. Fluorescently tagged *A. agrestis* β-CA1 confirmed the cytoplasmic localization (Supplementary Fig. 4). This absence of a diffuse stromal β-CA in *A. agrestis* is consistent with models of a pCCM ^13^ and reinforces that the relocalization of stromal CAs to cytoplasm is a key step toward engineering CCMs in crop plants ^30^.

### CAH3 localizes to the space between pyrenoids

While a stroma-localized CA would prohibit pCCM function, treatment with a CA inhibitor (ethoxzolamide) greatly decreased the rate of photosynthesis and significantly increased the CO_2_ compensation point of hornworts, but not liverworts ^31^, demonstrating that a CA is required for hornwort pCCM. In Chlamydomonas, CAH3 is located in thylakoid tubules, which form a knot in the center of the pyrenoid ^32–34^. Chlamydomonas CAH3 converts lumenal HCO_3_^-^ to CO_2_, allowing for diffusion of CO_2_ across the thylakoids and into the pyrenoid space. We discovered that the genomes of hornworts and some other bryophytes possess orthologs of the Chlamydomonas CAH3 gene, while vascular plants appear to have lost it (Fig. 3A; Supplementary Fig. 5). *A. agrestis* CAH3 is abundantly expressed, and retained not only the three critical histidine residues at the active site ^35^, but also the lumenal transit peptide (Supplementary Fig. 6, Supplementary Table 1). Importantly, we found that fluorescently tagged *A. agrestis* CAH3 formed puncta localized to the periphery of pyrenoids, concentrated adjacent (<500 nm) to the most interior side of the pyrenoids (Fig. 3b,c; Supplementary Fig. 6; see Movie S2 for Z-stack). CAH3 thus appears to be at the center of the chloroplast and in the space between pyrenoids. This localization pattern supports CAH3’s role in supplying CO_2_ to the surrounding pyrenoids in the hornwort pCCM.

**Figure 3.**
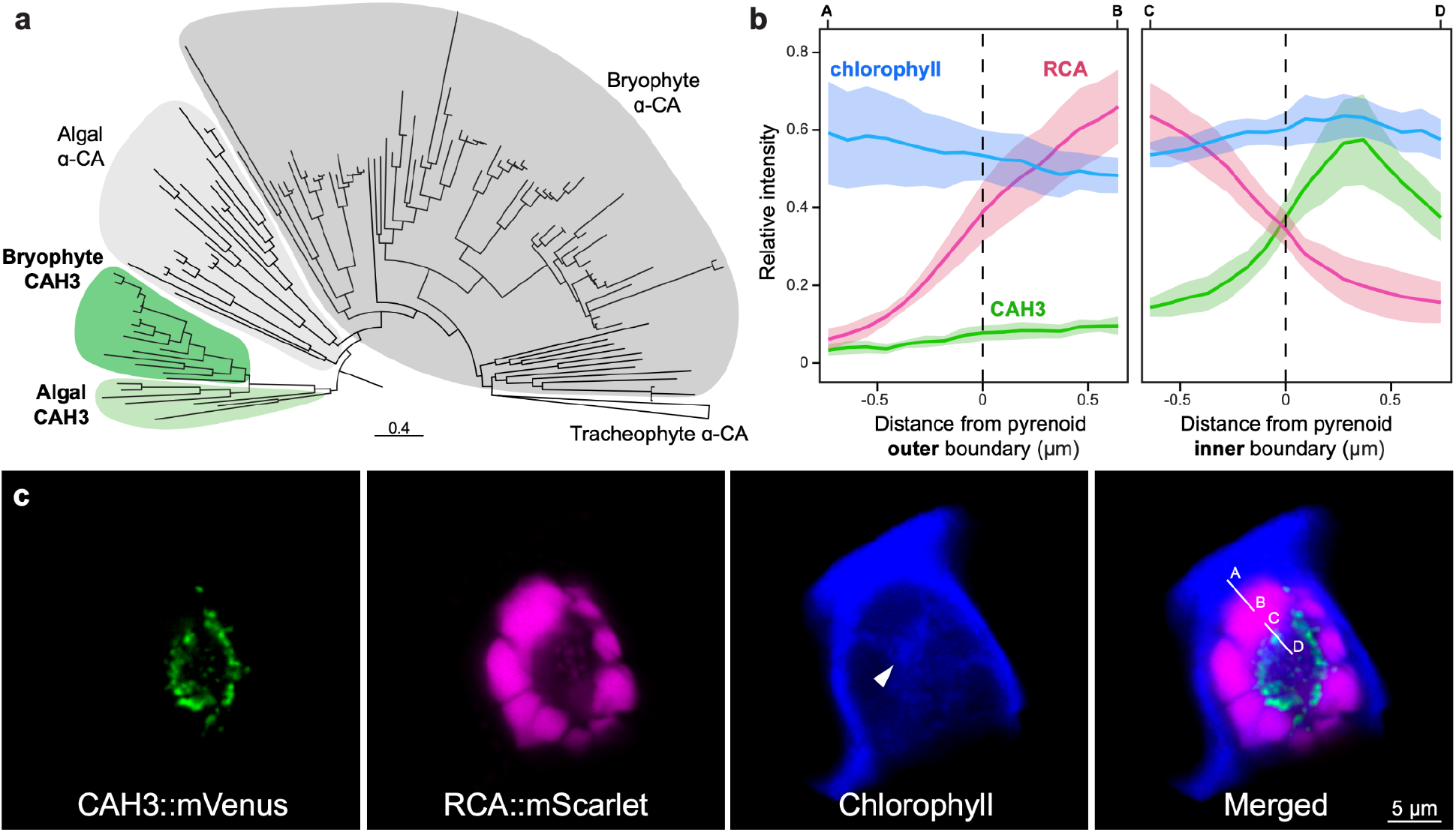
Pyrenoids are organized around CAH3 in *Anthoceros agrestis*. (a) Phylogeny of α-CA including CAH3. Orthologs of Chlamydomonas CAH3 can be found in hornworts and other bryophytes (shaded in green), but were lost in vascular plants. (b) CAH3::mVenus fluorescence intensity spikes at the interior side of pyrenoids (white lines in c). Error bars represent SEMs and are shown as shaded areas in each curve (n=10). (c) Example images of a cell co-transformed with CAH3::mVenus (green) and RCA::mScarlet (magenta). Chlorophyll autofluorescence is shown in blue. White arrowhead points to the central thylakoid knot. The Z-stack series can be found in Movie S2. (scale bar, 5 µm.)

### LCIB localizes to the chloroplast membrane

Another CA crucial to Chlamydomonas pCCM is LCIB ^36,37^, which localizes to the periphery of Chlamydomonas pyrenoids under low CO_2_ conditions to recapture leaked CO_2_ from pyrenoids^38^. LCIB orthologs are present in hornworts with conserved residues for coordination of Zn^2+^ ion and catalysis ^21^. Apart from hornworts, no LCIB homolog has been identified in other land plant genomes ^21^. This exclusive presence of LCIB in pyrenoid-bearing algae and hornworts suggests that LCIB may have a role in hornwort pCCMs. We discovered that, unlike LCIB in Chlamydomonas, fluorescently tagged *A. agrestis* LCIB did not localize to the immediate pyrenoid periphery, but instead to the edge of the chloroplast (Supplementary Fig. 7). To interrogate this pattern further, we co-localized LCIB::mVenus with mScarlet tagged TIC40, which is a known component of the inner chloroplast membrane ^39^. The fluorescence of LCIB::mVenus and TIC40::mScarlet clearly overlapped (Fig. 4), thus supporting that LCIB is indeed membrane-bound.

**Figure 4.**
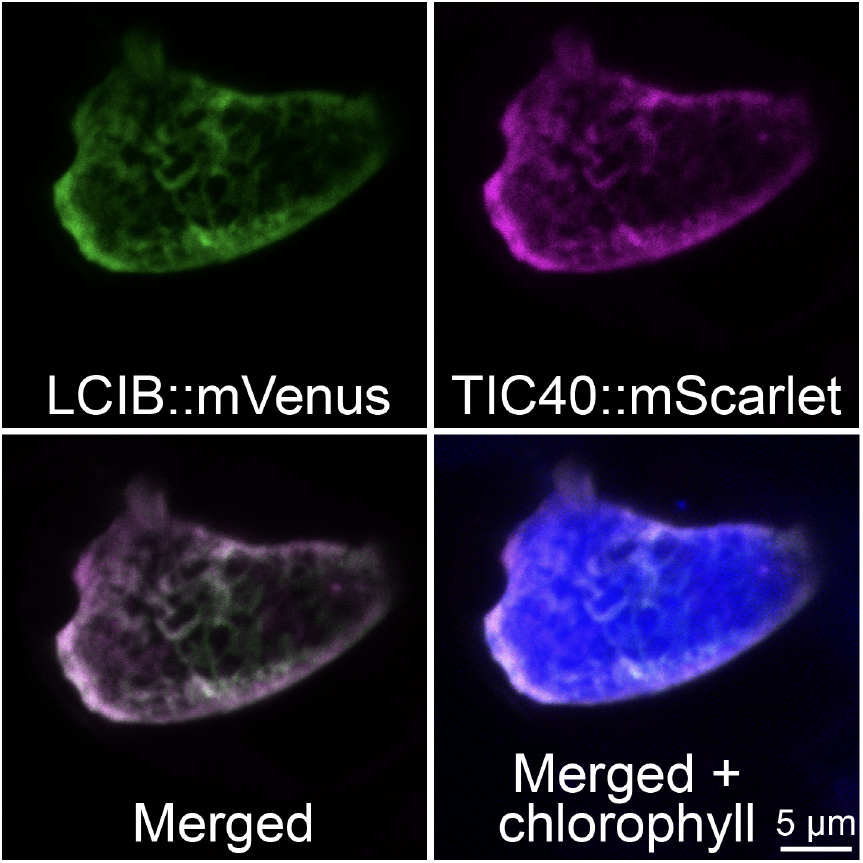
*A. agrestis* LCIB localizes to the chloroplast membrane. Maximum intensity projection of *A. agrestis* co-transformed with LCIB::mVenus (green) and TIC40::mScarlet (magenta). Chlorophyll autofluorescence is shown in blue. (scale bar, 5 µm.)

There are several reasons why the membrane-localized LCIB in *A. agrestis* is logical. In Chlamydomonas, LCIB localization to the starch sheath is likely mediated by its homolog LCIC^38^. Hornworts neither have starch sheaths nor LCIC, thus we would not expect to see a similar localization of LCIB. Furthermore, if LCIB were to surround each pyrenoid in hornworts, it would likely rehydrate any CO_2_ that had just been released by CAH3 (which are placed outside of pyrenoids; Fig. 3c), thus short-circuiting the pCCM. This scenario would be impossible in Chlamydomonas because CAH3 is localized in the thylakoid tubules embedded within a pyrenoid. Indeed, previous reaction-diffusion modeling of a pCCM showed that when thylakoid stacks are used as a diffusion barrier (as is the case of hornworts), LCIB localization to the chloroplast envelope would be optimal. This is because localizing LCIB as far from pyrenoids as possible minimizes “stealing” of CO_2_ from the pyrenoid matrix and can still serve as the last line of defense for CO_2_ leakage ^13^. Furthermore, for any passively diffused CO_2_ coming into the chloroplast, membrane-localized LCIB would quickly convert it into HCO_3_^-^ before it can escape back to the cytoplasm.

### Co-immunoprecipitation and genome-scanning did not reveal putative linker proteins

Pyrenoids, which house Rubisco, are central to pCCM. Rubisco linker proteins that mediate pyrenoid formation have been identified in the green algae Chlamydomonas (EPYC1) ^7^ and *Chlorella sorokiniana* (CsLinker) ^40^, as well as in the diatom *Phaeodactylum tricornutum* (PYCO1) ^25^. A putative linker was also proposed in the chlorarachniophyte alga *Amorphochlora amoebiformis* ^41^. While there is little sequence similarity between these linker proteins, they share certain physicochemical properties, and importantly, can all be pulled down using co-immunoprecipitation (co-IP) with an anti-Rubisco antibody. To identify the possible Rubisco linker in *A. agrestis*, we optimized the lysis of *A. agrestis* tissue and conducted co-IP using a custom anti-Rubisco antibody with soluble proteins clarified from hornwort lysate (Supplementary Fig. 8). From our proteomic analysis we found that both Rubisco large and small subunits, as well as RCA were highly enriched in the immunoprecipitated fraction relative to the control where no antibodies were applied (Supplementary Fig. 9). Gene ontology terms relating to chloroplast, photosynthesis, and thylakoid membranes were significantly overrepresented in the immunoprecipitated sample, likely reflecting the tight association between pyrenoids and thylakoids (Supplementary Fig. 10). Together, these data suggest that our co-IP approach was successful in enriching the bait (i.e. Rubisco) and its prey proteins (i.e. potential interactors).

Of the proteins significantly enriched by Rubisco co-IP, we searched for ones with similar physicochemical properties to other known Rubisco linker proteins. Namely, the candidate protein needs to: (1) be highly expressed, (2) be rich in repeat motif sequences, (3) be highly disordered, and (4) have a high isoelectric point (see Methods for detail). Only one protein was found (AnagrOXF.S1G403800.t1) possessing promising characteristics as a linker protein, which we named Putative Pyrenoid Protein 1 (PPP1). In addition to having an oscillating disorder profile, a chloroplast transit peptide, and high enrichment in the co-IP fraction, PPP1 has similarity to Structural Maintenance of Chromosomes (SMC) proteins (Supplementary Fig. 11). We found that fluorescently tagged PPP1 did not localize to the pyrenoid, but instead to the stroma (Supplementary Fig. 11). Interestingly, in some instances its localization became more concentrated in a region central to the chloroplast (Supplementary Fig. 11), reminiscent of the CAH3 localization pattern (Fig. 3). In any case, it is clear that PPP1 is not a Rubisco linker protein. Despite that our co-IP experiment found no evidence for a canonical linker in *A. agrestis*, we cannot rule out the possibility that one exists which has evaded our notice.

Therefore, we broadened our search to include the entire predicted proteome of *A. agrestis*, using the same screening criteria as before. Only one candidate protein, PPP2 (AnagrOXF.S4G428300.t2), was found meeting these criteria. Compared to the known Rubisco linkers (EPYC1, CsLinker, and PYCO1), the repeats of PPP2 were much more irregular and spaced erratically across the protein sequence (Supplementary Fig. 12). Using fluorescent protein tagging, we showed that PPC2 did not localize to the pyrenoid, but instead to the spaces between pyrenoid subunits (Supplementary Fig. 12). Similar to PPP1, no evidence supports PPP2 being a linker protein for *A. agrestis* Rubisco. Our results thus point to a possibility that *A. agrestis* might employ a different mode of pyrenoid condensation from what has been reported from algae.

### *A. agrestis* pyrenoids contain proteins involved in Rubisco assembly and Calvin-Benson-Bassham cycle

While the Rubisco linker in *A. agrestis* remains elusive, we found other proteins of interest that were enriched by immunoprecipitation. For example, Rubisco assembly factors (Raf1, Cpn60α, Cpn60β) showed significant enrichment by co-IP (Supplementary Fig. 9). We tested the localization of Cpn60β, a chaperonin required for Rubisco folding ^42,43^ by fluorescent protein-tagging, and found Cpn60β::mVenus to be clearly pyrenoid-localized (Fig. 5a, Supplementary Fig 13). To further explore if Rubisco biogenesis might occur within pyrenoids, we tagged another Rubisco assembly factor, RbcX2 ^44^. While RbcX2 was not co-immunoprecipitated, it is also localized to the pyrenoids (Fig. 5b, Supplementary Fig 13).

**Figure 5.**
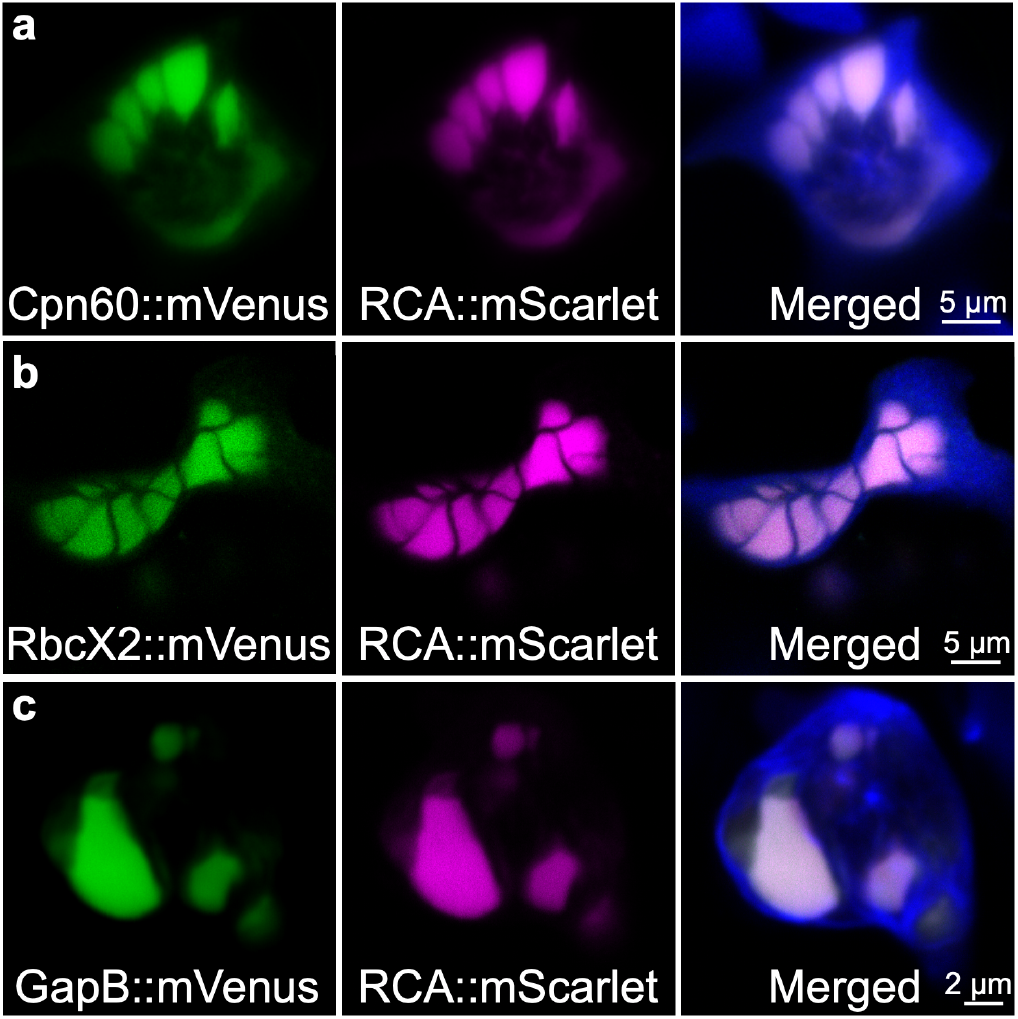
*A. agrestis* Rubisco assembly and CCB cycle proteins localize to pyrenoids. Example images of cells expressing either (a) Cpn60::mVenus, (b) RbcX2::mVenus, or (c) GapB::mVenus tagged with mVenus (green). RCA::mScarlet was co-transformed to mark pyrenoids (magenta). The merged images also included chlorophyll autofluorescence (blue). (scale bar, 2 or 5 µm.)

Another protein enriched by co-IP was glyceraldehyde-3-phosphate dehydrogenase (GAPDH). Chloroplastic GAPDH catalyzes the reductive step of the Calvin-Benson-Bassham (CBB) cycle and is composed of A and B subunits, both of which were significantly enriched in the immunoprecipitated fraction (Supplementary Fig. 9). To examine if GAPDH is a pyrenoid component, we fluorescently tagged the B subunit (GapB) and confirmed its pyrenoid localization (Fig. 5C, Supplementary Fig 14). It is possible that the CBB cycle (or part of it) takes place in pyrenoids to limit the diffusion time of CBB cycle intermediates. Contrastingly, in Chlamydomonas, CBB enzymes are localized to the periphery of the pyrenoid ^45^.

Based on our results, it is plausible that *A. agrestis* pyrenoids are an inclusive compartment, where multiple cellular processes revolving around Rubisco take place. Future studies systematically tagging Rubisco assembly and CBB cycle proteins are needed to further test this hypothesis.

## DISCUSSIONS

### A working model for a hornwort pCCM

Our current evidence suggests a model (Fig. 6) for the hornwort pCCM that bears similarity to the algal pCCM yet with some key differences. First, since hornworts largely do not exist in an aqueous environment (where CO_2_ diffusion is severely limited), we suspect that periplasmic CAs and active HCO_3_^-^ transporters such as LCI1/HLA are not essential. Likewise, despite retaining the LCIB-BST-CAH3 chassis, hornworts do not have an ortholog to LCIA, the hypothesized HCO_3_^-^ channel in Chlamydomonas. Given this, we hypothesize that the hornwort CCM likely operates under a “passive”, LCIB-dependent mode as described by Fei et al ^13^.

**Figure 6.**
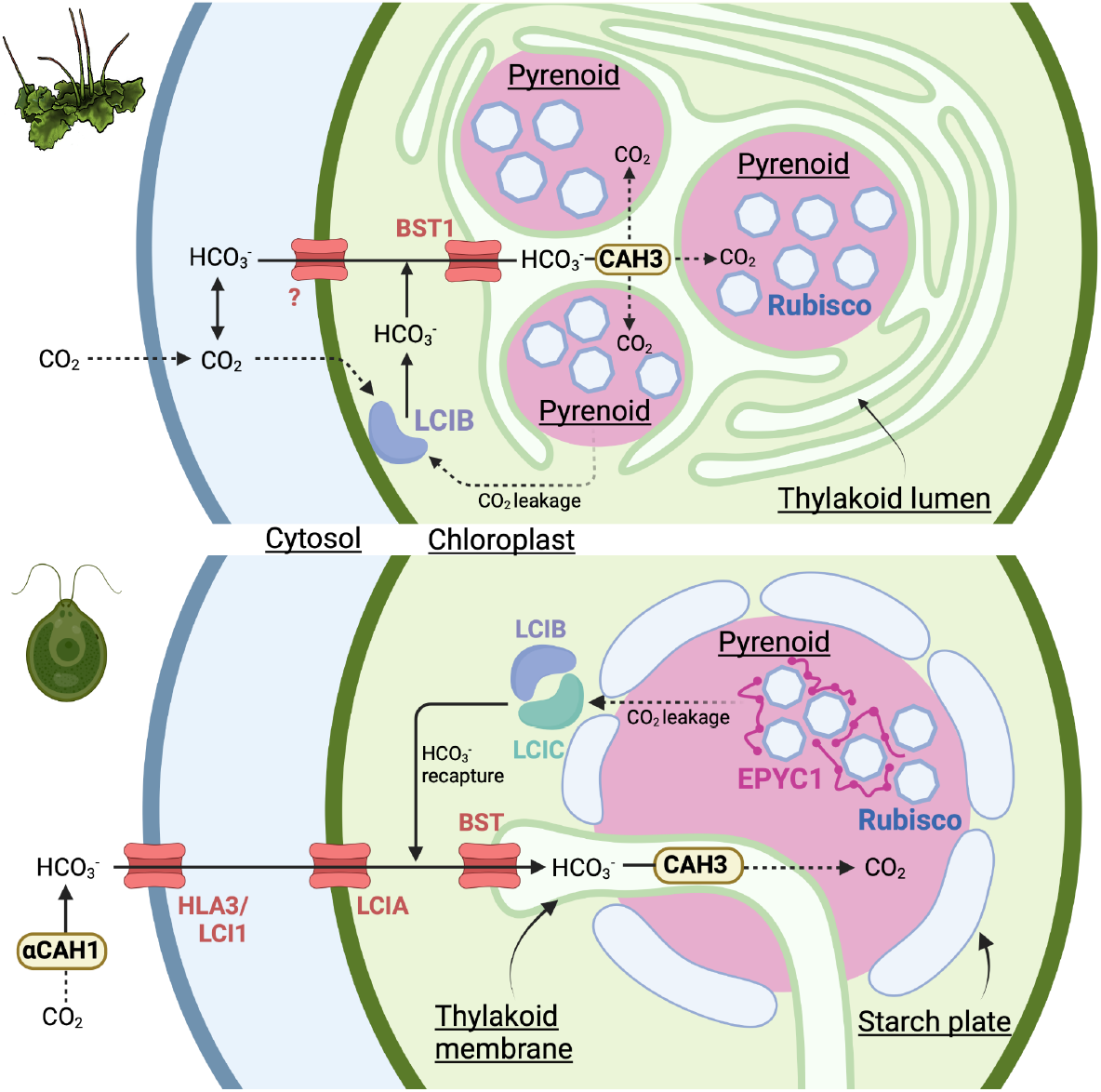
The spatial model of *A. agrestis* pCCM and (top) comparison with that of Chlamydomonas (bottom). The hornwort illustration was done by Rebecca Key.

Once CO_2_ passively diffuses across the chloroplast membrane it is rapidly converted to HCO_3_^-^ by membrane-localized LCIB, trapping it in the stroma. In the stroma, HCO_3_^-^ can enter the thylakoid space via BST1, where a concentration gradient drives it towards the pyrenoid adjacent thylakoids enriched in CAH3. There, CAH3 catalyzes the dehydration of HCO_3_^-^, creating a CO_2_ enriched environment for the nearby pyrenoids. Leakage of CO_2_ from the pyrenoid matrix is hindered by the highly reticulated thylakoid network, and any escaped CO_2_ could be recaptured by LCIB at the chloroplast membrane. While we cannot rule out the presence of active Ci transport, the condition to drive an efficient pCCM is met based on the localizations of LCIB, BST, and CAH3, thylakoid-enclosed pyrenoids, as well as the reaction-diffusion model by Fei et al ^13^.

The spatial model we presented here provides a framework for future hypothesis-driven research. Although protein localization alone does not provide the full proof for a protein’s function, currently no reverse genetics method has been successfully developed in *A. agrestis*. Future work investigating carbon assimilation efficiency of selective gene knockdown or knockout mutants, as well as biochemical *in vitro* characterizations will be crucial to confirm or revise our model.

### The first land plants likely had the chassis for a biophysical CCM

Operating an effective biophysical CCM requires the strategic placement of Ci channels and CAs. It is striking that some of these key components (LCIB, BST, and CAH3) characterized in Chlamydomonas have orthologs in hornworts and likely play similar roles in their respective pCCM. It is difficult, if not impossible, to determine whether hornwort pCCMs represent an ancestral trait or an independent origin. In either case, however, the presence of the LCIB-BST-CAH3 chassis in hornworts suggests that the underlying chassis to concentrate CO_2_ was present in the last common ancestor of land plants. Considering that various algae possess biophysical CCMs even in the absence of pyrenoids ^46–49^, it is not impossible that the earliest land plants could have operated a functional CCM.

We propose two scenarios that could result in hornworts being the only land plants having a pCCM. In one, pyrenoids were *de novo* evolved in hornworts on top of the LCIB-BST-CAH3 chassis already in place. The loss of LCIB and CAH3 in other plant lineages rendered the evolution of pyrenoids less advantageous. The supposed absence of a canonical Rubisco linker for pyrenoid formation in *A. agrestis* lends support for the separate origins of pyrenoids. Alternatively, pyrenoids could be ancestral to land plants, but were subsequently lost once in setaphytes (mosses and liverworts) and another time in vascular plants. In this scenario hornworts did not independently evolve pyrenoids, but rather retained them as a plesiomorphic trait.

### Implications for the repeated evolution of pyrenoids within hornworts

Within hornworts, pyrenoids have been lost and gained multiple times across the phylogeny ^50^. Interestingly, LCIB, BST, and CAH3are present in all the sequenced hornwort species, even in those lacking pyrenoids ^51^. Considering that pyrenoid-absent species have CO_2_ compensation points which are intermediate between liverworts and pyrenoid-containing hornworts ^52^, a baseline biophysical CCM probably exists among most hornworts and the strength of this CCM could be accentuated by the presence of a pyrenoid. The universal presence of this LCIB-BST-CAH3 chassis could explain the repeated gains of pyrenoids as it might have predisposed the evolution of pyrenoids. The specific localization of CAs and Ci channels is also likely the first step to the evolution of a fully functional pCCM, rather than pyrenoid formation itself.

### Hornworts provide a blueprint for engineering a biophysical CCM in land plants

Significant strides have been made in transplanting parts of the algal CCM to Arabidopsis in recent years. For example, by introducing EPYC1, SAGA1, and SAGA2 proto-pyrenoids with starch sheaths have been successfully built in Arabidopsis ^53,54^. However, the entire algal CCM module has yet to be integrated into flowering plants. Our investigation into the hornwort system demonstrates that some components of the algal pCCM may not be required in land plants.

First, a starch sheath around pyrenoids may not be necessary as thylakoid stacks could provide a sufficient CO_2_ diffusion barrier (Fig. 1). Second, if hornworts indeed operate a “passive” pCCM, then our work provides the first empirical evidence for its feasibility ^13^. In a passive pCCM, the proper localization of a minimal set of carbonic anhydrases (e.g. LCIB and CAH3) and thylakoid localized Ci channels (e.g. BST) may be sufficient to concentrate CO_2_ without the need for active Ci transport to the chloroplast ^13^. Importantly, recent modeling suggested that introducing active HCO_3_^-^ transporters to build pCCMs in land plants is only viable if chloroplast membrane permeability to CO_2_ is low ^55^. The passive pCCM of hornworts would not encounter this challenge and could be potentially simpler to engineer. Taken together, hornworts provide an alternative blueprint to design a future pCCM.

## METHODS

### *Anthoceros agrestis* cultures

*Anthoceros agrestis* axenic cultures were maintained on *A. agrestis* gametophyte growth medium (AG medium ^56^) supplemented with 0.2% sucrose following the protocol described in Lafferty et al ^23^. The growth conditions included a temperature of 22°C, a 16/8 h light/dark cycle, and light intensity ranging from 6 to 25 μmol/m^2^/s. Warm white and cool white light was emitted by Ecolux XL Starcoat F32T8 XL SP30 ECO fluorescent bulbs and F32T8 XL SP41 ECO bulbs, respectively (General Electric, USA). A total of 3-6 g of fresh thallus tissue was homogenized and prepared for transformation as described in Lafferty et al ^23^. The homogenate was filtered using a 100 μM cell strainer (MTC Bio, USA), and the thallus tissue was washed with sterile deionized water and plated on AG medium supplemented with 2% sucrose.

### Transmission electron microscopy

Plant materials were processed for transmission electron microscopy following the published protocol ^57^, with some modifications. Pure cultures were fixed in 3% glutaraldehyde, 1% formaldehyde, and 0.75% tannic acid in 0.05 M Na-cacodylate buffer, pH 7, for 3-4 h at room temperature. After several rinses in Sorensen’s 0.1 M buffer, the samples were post-fixed in 2% osmium tetroxide, dehydrated in an ethanol series and embedded in Spurr’s resin for 24 h. Thin sections were cut with a diamond knife, stained with methanolic uranyl acetate for 15 min followed by Reynolds’ lead citrate for 10 min, and observed with a Hitachi H-7100 transmission electron microscope at the Imaging-Microscopy Platform of the IBIS, Université Laval.

### Construction of fluorescently tagged protein expression cassettes

Constructs were designed and built using the OpenPlant toolkit ^58^. All genes of interest were driven by the same constitutive promoter (*A. agrestis* native *Elongation factor 1 alpha* (*Ef1α*) promoter) and *Nos* terminator (OP-053). Coding sequences of target genes were synthesized by Twist Biosciences with synonymous substitutions made where necessary to remove internal *Bsa*I or *Sap*I Type II restriction cut sites or to reduce repetitive nucleotide sequences if required for synthesis. Assembly reactions were performed by a one-pot mixture that consists of Type II restriction enzymes with T4 DNA ligase (NEB, USA) as described in Sauret-Gueto et al. ^58^. Heat shock transformation with NEB® 5-alpha Competent *Escherichia coli* was used to regenerate ligated products on the respective Luria Broth agar plates with either kanamycin or streptomycin. Colony PCR was performed with KOD One™ PCR Master Mix (Toyobo) to screen colonies before recovering plasmids with a 5 mL overnight culture and miniprep (QIAgen).

Whole plasmid sequencing (Eurofin Genomics, USA) was performed as a final validation. To prepare plasmids for biolistic mediated transformation, a total of 50 ml *E. coli* culture was grown overnight at 37 °C. The plasmid DNA was extracted using the PureYield™ Plasmid Midiprep system (Promega, USA), followed by vacuum concentration in a SpeedVac Concentrator (Thermofisher Scientific, USA) to obtain DNA concentrations of 1 μg/μL.

### Transient expression of fluorescently tagged proteins in *A. agrestis*

To obtain transient expression of fluorescently tagged proteins in *A. agrestis*, we performed biolistic mediated transformation using DNA/gold particle mixture preparations and the particle Delivery system PDS-1000/He (Bio-Rad, USA) as described in Lafferty et al ^23^, with a target distance of 14 cm and a burst pressure of 450 psi, under a vacuum of 28 inHg. The bombarded tissue was left to recover for 3-10 days, under standard culturing conditions as described above, before imaging.

### Confocal imaging

Imaging was performed using Zeiss LSM710, Leica TCS SP5 Laser Scanning Confocal Microscope, or Leica Stellaris 5 confocal microscope. Proteins tagged with mVenus were excited with a 514 nm laser line with an emission band of 524/551 nm. Chlorophyll autofluorescence and proteins tagged with mScarlet were excited with a 561 nm laser line, with emission bands of 658/699 and 579/609 nm, respectively. Each FRAP experiment started with four initial scans before bleaching the region of interest (ROI). Circular ROIs with a 1.3 µm diameter were exposed eight times at 20% intensity of the 514 nm laser, before allowing recovery for 160 cycles (66 seconds) with laser power attenuated at 0.5% intensity. Replicates were collected by photobleaching individual pyrenoids within a cell. Subsequent analysis was conducted using the FRAP wizard in Leica Application Suite Advanced Fluorescence and data was plotted in R. Intensity profiles for all tagged proteins were collected by using the line tool to transect regions of interest (e.g. pyrenoid boundaries, chloroplast boundaries) and then plotting the intensity profile in Fiji ^59^. These intensity profiles were then exported for subsequent analyses and plot generation in R.

### Homology-based approach to identify candidate CCM proteins

To identify candidate CCM genes in hornworts we used Orthofinder v2.5.4 on a broad sampling of plant and algal genomes (Supplemental Table 2), including 11 species of hornworts ^51^, to generate orthogroups. We then selected orthogroups containing key Chlamydomonas CCM genes (Supplemental Table 1). Because orthogroups can sometimes represent large gene families, to infer the direct orthologues in hornworts, phylogenetic reconstruction using IQ-TREE ^60^ was performed to identify the hornwort sequences in the same subfamily as the Chlamydomonas genes of interest. From these, we selected *A. agrestis* genes which showed high expression levels using the RNA-seq data from Li et al ^21^. In cases where there were multiple orthologues present in *A. agrestis*, we selected the gene with the highest expression. In addition to CCM genes, we choose Photosystem I reaction center subunit VI (PSAH) and Oxygen-evolving enhancer protein 2 (PSBP) as Photosystem I and Photosystem II markers, respectively, to compare the distribution of photosystems and to visualize thylakoids.

### Targeted bioinformatic search for a Rubisco linker protein

A targeted search for potential linker proteins was conducted in a manner similar to, but less stringent than described in Mackinder et al ^7^. Genomic sequences of two *A. agrestis* strains ^21^ were first screened for tandem repeats using Xstream ^61^. Default parameters were used with the following exceptions: Minimum Period: 20; Maximum Period: 100; Minimum Copy # 2; and Minimum tandem repeat Domain: 40. Proteins which passed this screen were then analyzed for: 1) a high isoelectric point (>8) using Expasy ^62^, 2) a high proportion of disordered sequence using the PONDR server (>50%) ^63^. From this screen, lowly expressed genes were filtered based upon their corresponding RNA transcript abundance (transcript per million <10) ^21^.

### Optimizing lysis of *A. agrestis* thallus

To derive soluble proteins from *A. agrestis* thallus, we experimented the lysis of *A. agrestis* by testing four different buffers: buffer A (20 mM HEPES pH 7.0, 50 mM NaCl, 12.5% v/v glycerol), buffer B (Buffer A with 2% v/v Triton X-100), buffer C (20 mM CAPS pH 11.0, 50 mM NaCl, 12.5% v/v glycerol), and buffer D (Buffer C supplemented with 2% v/v Triton X-100). For each lysis, 5 g of *A. agrestis* tissue were harvested from liquid cultures in the AG medium ^56^. The biomass was frozen with liquid N_2_ and crushed to find powder with a mortar and pestle. A total of 5 mL of either test buffer supplemented with 1 mM phenylmethylsulfonyl fluoride and one protease inhibitor tablet (Roche) was then to resuspend the fine powder. The suspension was lysed with a French pressure cell press (American Instrument Company, USA) at 1000 PSI. Polyvinylpolypyrrolidone (PVPP, 2%) was added to the French pressed lysate and the mixture was sieved through four layers of Miracloth, pre-wetted with the respective test buffer. The filtrate was centrifuged (21,000 *g* for 45 min at 4 °C) to separate pellet and supernatant fractions. The supernatant fraction was further filtered through a 0.22 µm syringe filter. Both fractions were analyzed on SDS-PAGE and western blot using 8–16% Mini-PROTEAN® TGX™ Precast Protein Gels (Bio-Rad). For immunoblotting of Rubisco, a polyclonal antibody was raised in rabbits against the *A. agrestis* Rubisco large subunit C-terminus peptide EVWKEIKFVFETIDTL and affinity purified (Life Technologies, USA).

### Co-immunoprecipitation of Rubisco

A total of 100 µL Protein A resin (Dynabeads, Invitrogen) was aliquoted into 1.5 mL microcentrifuge tubes and washed thrice with buffer C (20 mM CAPS pH 11.0, 50 mM NaCl, 12.5% v/v glycerol) with a magnetic rack. Next, the resin was incubated with 7.5 µg/mL anti-Rubisco on a rotator (2 h/4°C) and washed twice with buffer C. Controls were performed by not priming the resin with anti-Rubisco antibodies. A total of 750 µL soluble lysate was then added to anti-Rubisco bound resin and incubated on a rotator overnight (∼18 h/4 °C) and washed thrice with buffer C. Protein elution was carried out by adding 80 µL of 2.5X SDS loading buffer, without 2-Mercaptoethanol. Eluted proteins were separated from the Dynabeads and 2-Mercaptoethanol was added to a final concentration of 100 mM, proteins were then boiled (95 °C/5 min) and shipped for LC-MS/MS analysis. The co-IP and control experiments each had five technical replicates.

### Proteomic analysis

LC-MS/MS analysis was performed at the Environmental Molecular Sciences Laboratory in Pacific Northwest National Laboratory. Samples were processed using Filter Aided Sample Preparation (FASP) ^64^ by adding 400 µl of 8 M urea to 30K molecular weight cut off (MWCO) FASP spin columns along with 40 µl of the sample in SDS BME buffer and centrifuged at 14,000 *g* for 20 min. Urea washes were repeated three additional times followed by the addition 400 µl of 50 mM ammonium bicarbonate, pH 8.0 and two repeated centrifugation for 20 min. The columns were then placed into clean and labeled collection tubes. The digestion solution was made by dissolving 5 μg trypsin in 75 μL 50 mM ammonium bicarbonate solution which was added to each sample. The samples were then incubated for 3 h at 37 °C with 600 rpm shaking on a thermomixer with a ThermoTop (Eppendorf, Hamburg, Germany) to reduce condensation into the caps of collection tubes. The resultant peptides were then centrifuged through the filter and into the collection tube by centrifuging at 14,000 *g* for 15 mins. The peptides were concentrated to ∼30 µL using a vacuum concentrator. Final peptide concentrations were determined using a bicinchoninic acid (BCA) assay (Thermo Scientific, Waltham, MA USA) and each sample was prepared at 0.1 µg/µl for MS analysis Digested protein samples were analyzed using an Orbitrap Eclipse Tribrid MS (Thermo Scientific) outfitted with a high field asymmetric waveform ion mobility spectrometry (FAIMS) interface, using data-dependent acquisition mode. Peptides were ionized using a voltage of 2.4 kV and with an ion transfer tube temperature at 300 **°**C. Data acquisition time was 2 hr following a 20 min delay to avoid dead time between injection and elution of peptides. A proprietary method for transferring identification based on FAIMS filtering was used to fractionate ionized peptides by the FAIMSpro interface using a 3-Compensation Voltage (3-CV); -45, -60, -75 V method. Fractionated ions with a mass range 400-1800 m/z were scanned with Orbitrap at 120,000 resolution with an injection time (IT) of 50 ms and an automatic gain control (AGC) target of 4e5. Cycle times of 1.0 s were used for the 3-CV method. Precursor ions with intensities > 1e4 were fragmented with an isolation window of 0.7 by 30% higher-energy collisional dissociation energy and scanned with an AGC target of 1e^4^ as well as an IT of 35 ms.

Raw data files were referenced to *A. agrestis* nuclear encoded and chloroplast encoded proteins ^21^, and peptide abundances were extracted from the raw spectra using MASIC ^65^ and log2 transformed to remove skewness in distribution of measured abundances. Transformed abundance values were then normalized using the mean central tendency method implemented in InfernoR ^66^. Normalized peptide abundances were de-logged, summed, transformed (log2), and normalized again in InfernoR to produce normalized abundances for the protein level roll-up. Proteins which were missing in more than one replicate of the conditions (control or anti-RbcL) were filtered from the final analysis to limit the imputation of too many missing values.

Left-censored missing values were imputed using the Minimum Probability method with the default parameters. Differential enrichment analysis was conducted using the DEP Bioconductor package version 1.27.0 ^67^ with a p value cutoff of 0.05. Plots were generated using ggplot2 (version 3.5.1). Gene ontology (GO) enrichment analysis was carried out using the GO Enrichment module of TBtools ^68^ with *goslim_plant* selected. Background file was set as the entire *A. agrestis* proteome and proteins found to be significantly enriched by coimmunoprecipitation were chosen as the selection set. The resulting table was then used to generate an enrichment barplot.

## ACKNOWLEDGEMENTS

This work is supported by National Science Foundation MCB-2213841 to F.-W.L. and MCB-2213840 to L.H.G, Environmental Molecular Sciences Laboratory User Grant to F.-W.L., Triad Foundation Grant to F.-W.L., and Schmittau-Novak Graduate Student Grant to T.A.R. We thank Aleksandra Skirycz, Kathryn Eshenour, Amber Hotto, and David Stern for providing access to tools and reagents to establish preliminary lysis and co-IP experiments, Rebecca Key for the hornwort illustration, Carrie Nicora and Nikola Tolic from Environmental Molecular Sciences Laboratory in Pacific Northwest National Laboratory for technical assistance with mass spectrometry processing, the Plant Cell Imaging Center at Boyce Thompson Institute and Mamta Srivastava for technical assistance to confocal imaging with Leica TCS SP5 Laser Scanning Confocal Microscope, Adrienne Roeder for providing access to Zeiss LSM710 and Leica Stellaris 5 confocal microscope, the York Physics of Pyrenoids research community for feedback, and Li and Gunn lab members for discussions.

## AUTHOR CONTRIBUTIONS

T.A.R., L.H.G., and F.-W.L conceived the project. T.A.R. made the gene constructs and carried out confocal imaging. T.A.R., D.L., and X.X. performed hornwort transformation. Z.G.O. designed the Rubisco antibody and optimized the lysis of hornwort thallus. T.A.R. and Z.G.O. performed co-immunoprecipitation. J.C.A.V. performed transmission electron microscopy.

T.A.R., Z.G.O., L.H.G. and F.-W.L. analyzed the data. T.A.R. and F.-W.L. wrote the manuscript with contributions and comments from all authors. L.H.G. and F.-W.L. secured the funding and supervised the project.

