## Supplementary Figures and Tables for "Hornworts reveal a spatial model for pyrenoid-based CO_2_-concentrating mechanisms in land plants"

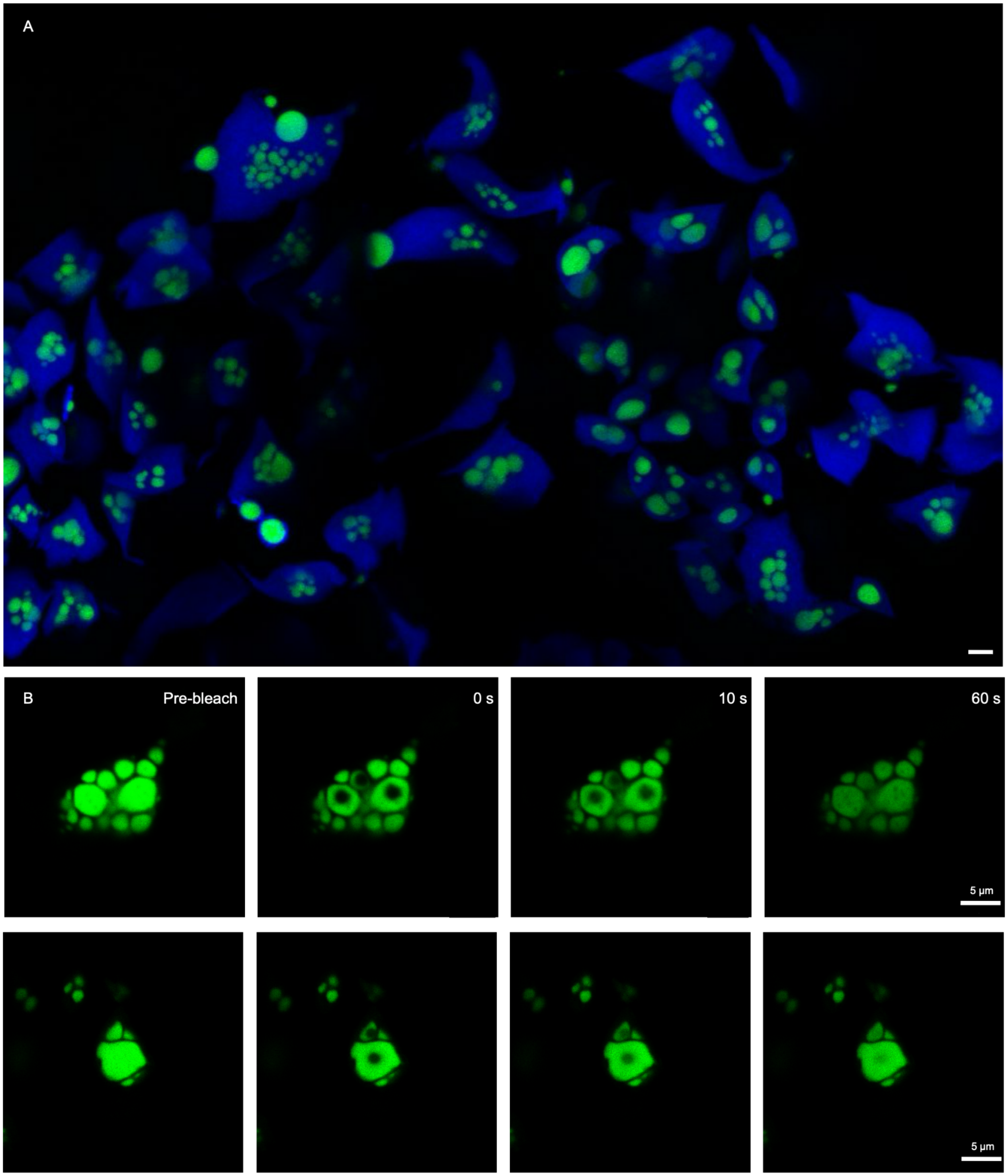


**Supplementary Fig. 1. Stable transgenic *A. agrestis* expressing RCA::mVenus**. (A) Pyrenoids of variable number and size were observed. Chlorophyll autofluorescence is shown in blue and RCA::mVenus fluorescence in green. (scale bar, 5 µm.) (B) Additional images of photobleached hornwort pyrenoids labeled with RCA::mVenus. (scale bar, 5 µm.)

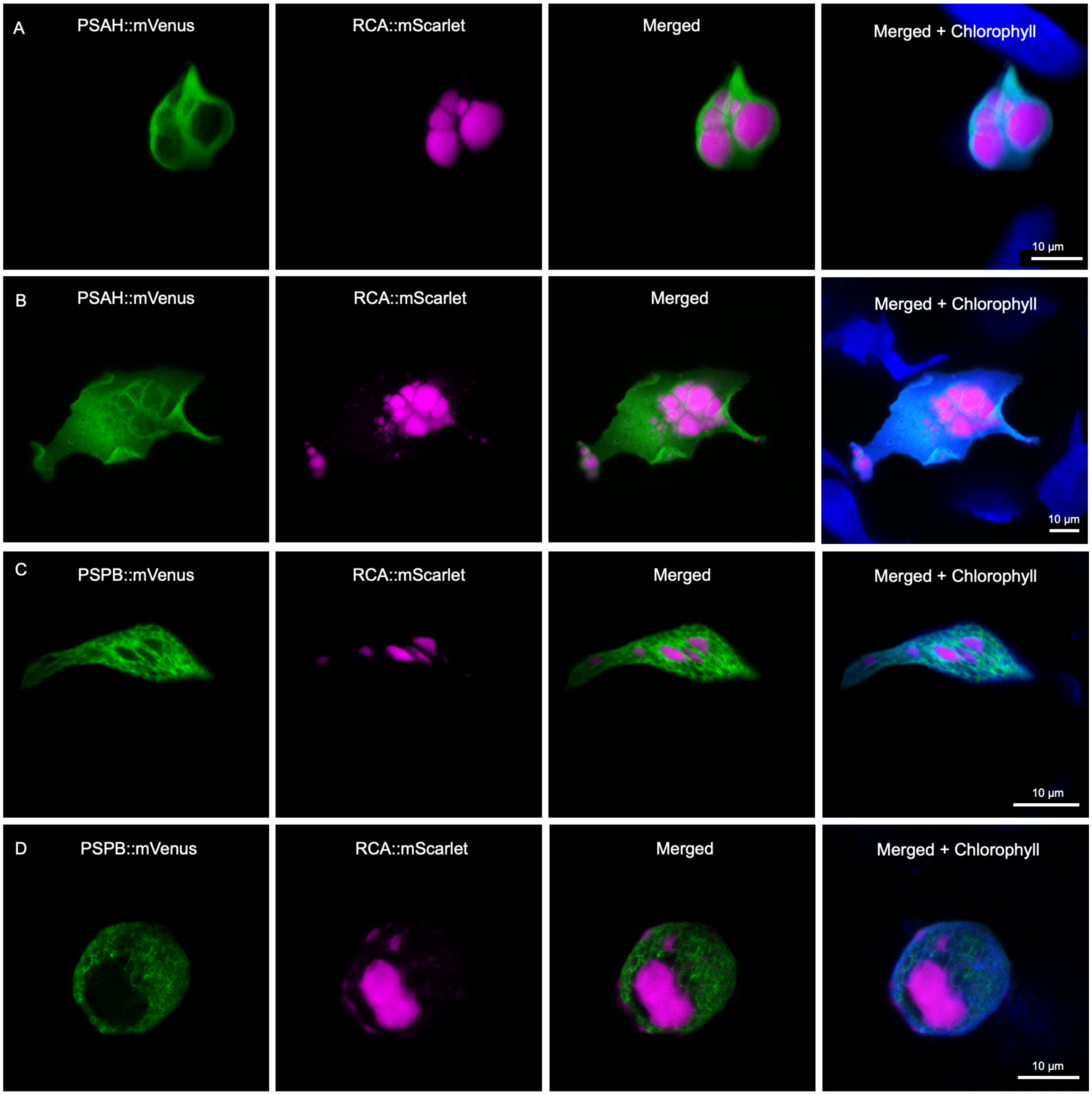


**Supplementary Fig. 2. Distribution of photosystems on thylakoids of *A. agrestis*.** Additional images of *A. agrestis* expressing (A and B) PSAH::mVenus and (C and D) PSBP::mVenus. mVenus tagged proteins are shown in green, RCA::mScarlet in magenta, and chlorophyll autofluorescence in blue. A and B are displayed as maximum intensity projections. (scale bar, 10 μm.)

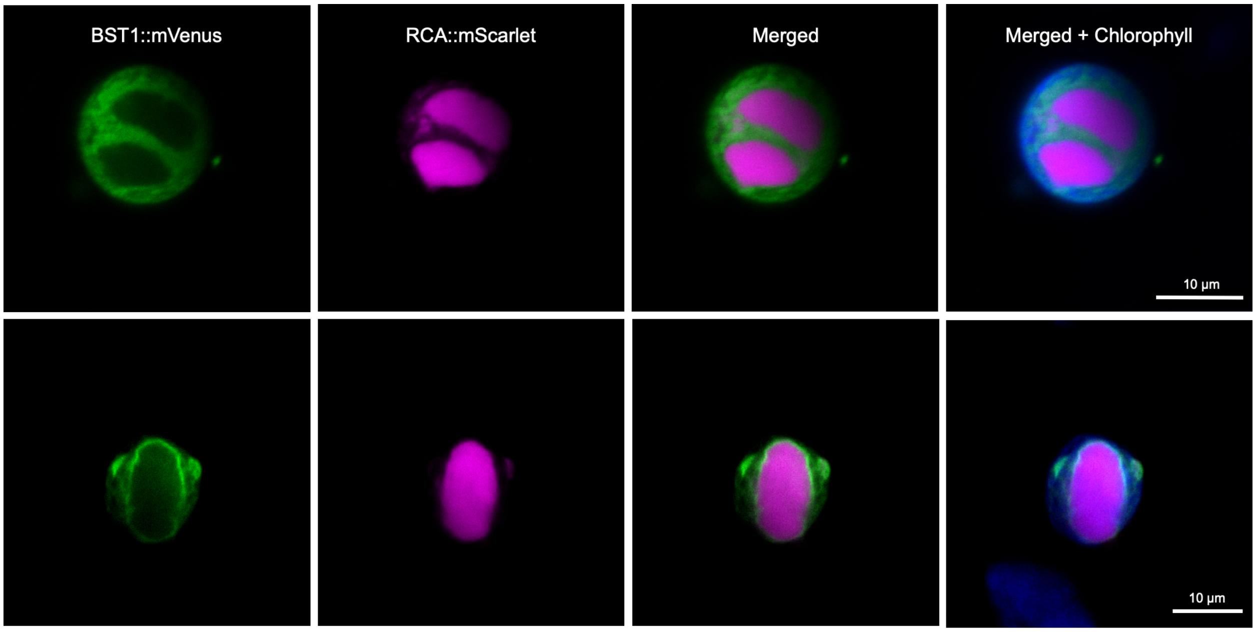


**Supplementary Fig. 3. Distribution of BST channels on thylakoids of *A. agrestis*.** Additional images of *A. agrestis* expressing BST::mVenus. mVenus tagged proteins are shown in green, RCA::mScarlet in magenta, and chlorophyll autofluorescence in blue. (scale bar, 10 µm.)

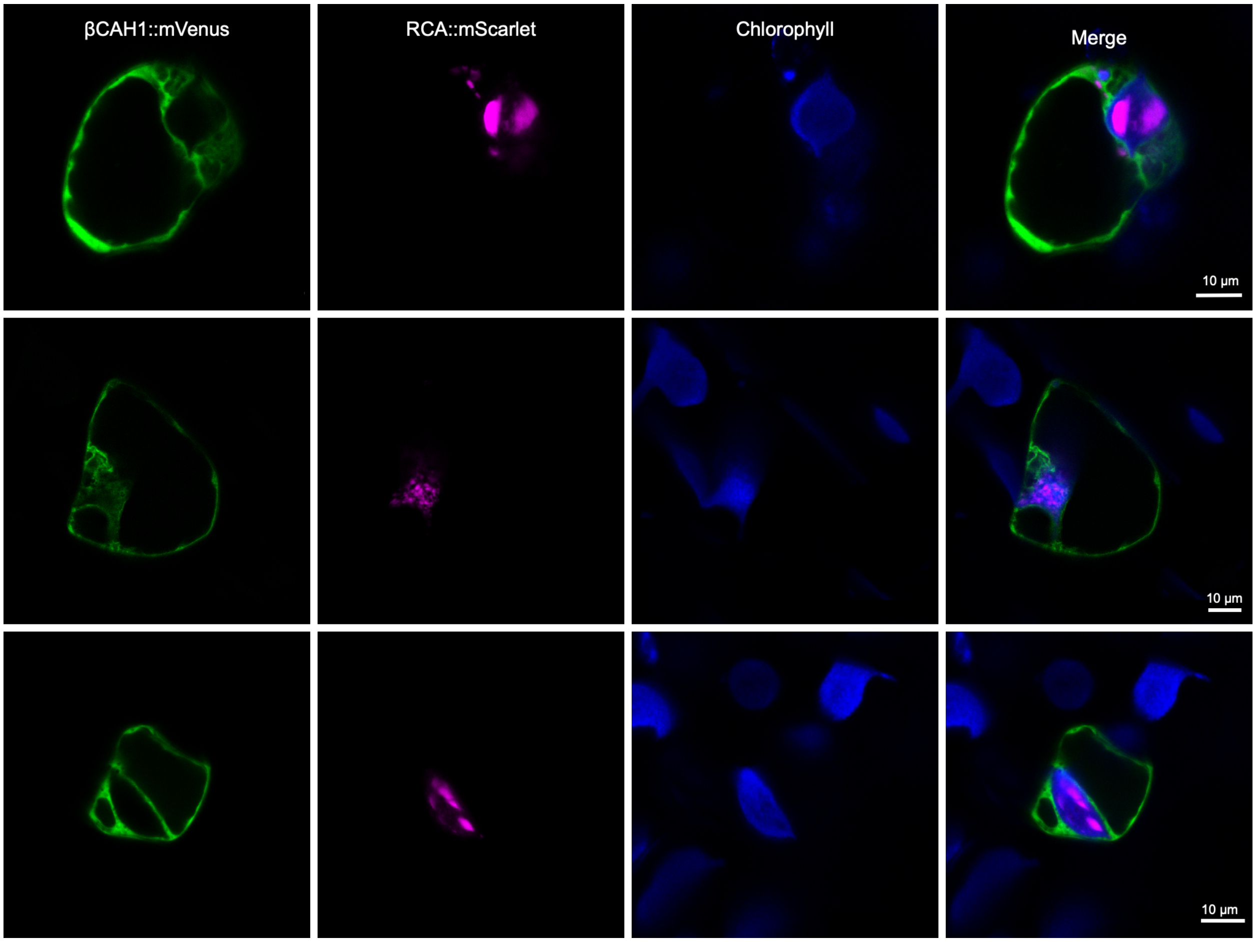


**Supplementary Fig. 4. β-CA1::mVenus localizes to cytosol in *A. agrestis*.** β-CA1::mVenus is shown in green, RCA::mScarlet in magenta, and chlorophyll autofluorescence in blue. (scale bar, 10 μm.)

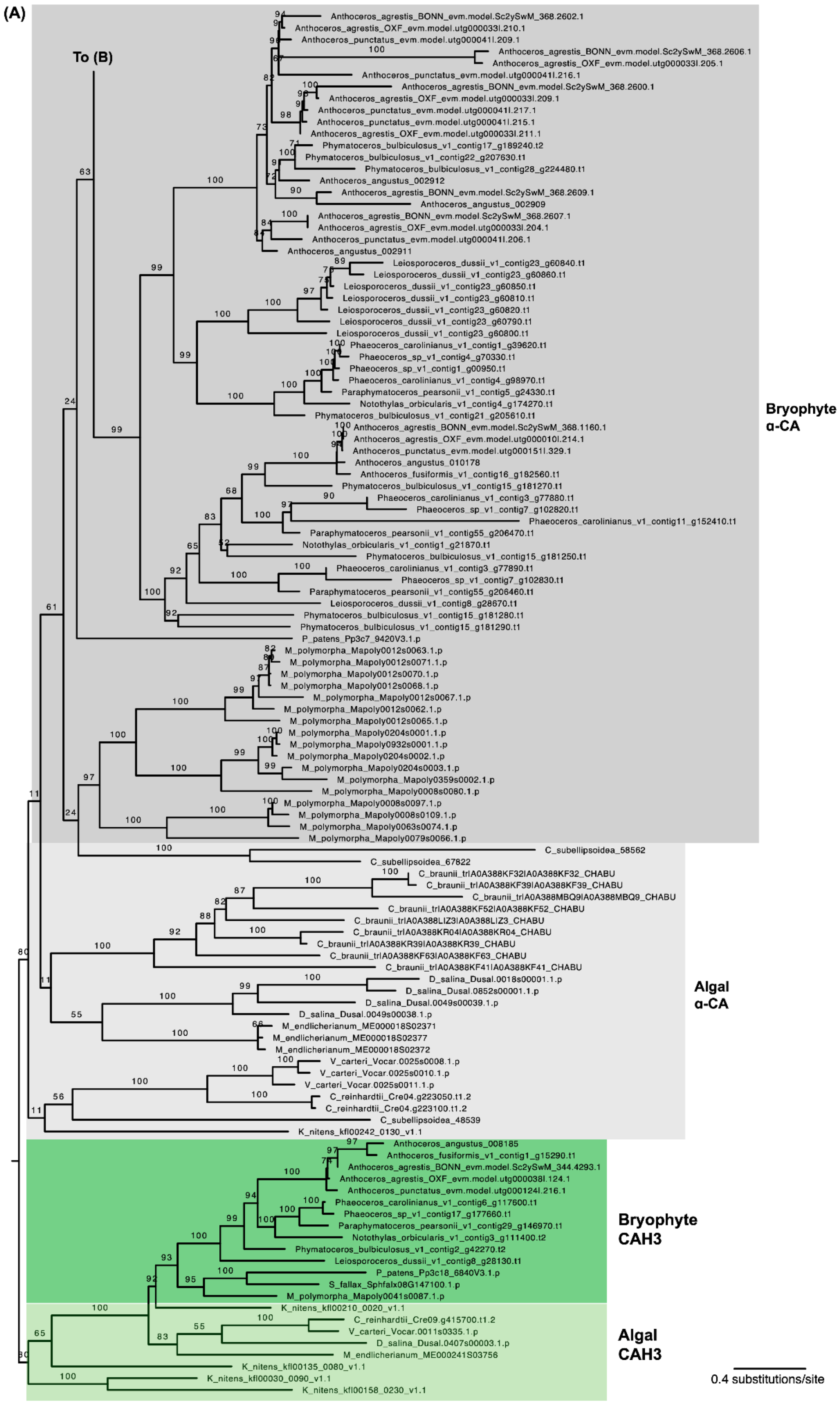

(A)
To (B)
Anthoceros_agrestis_BONN_evm.model.Sc2ySwM_368.2602.1
Anthoceros_agrestis_OXF_evm.model.utg000033l.210.1
Anthoceros_punctatus_evm.model.utg000041l.209.1
Anthoceros_agrestis_BONN_evm.model.Sc2ySwM_368.2606.1
Anthoceros_agrestis_OXF_evm.model.utg000033l.205.1
Anthoceros_punctatus_evm.model.utg000041l.216.1
Anthoceros_agrestis_BONN_evm.model.Sc2ySwM_368.2600.1
Anthoceros_agrestis_OXF_evm.model.utg000033l.209.1
Anthoceros_punctatus_evm.model.utg000041l.217.1
Anthoceros_punctatus_evm.model.utg000041l.215.1
Anthoceros_agrestis_OXF_evm.model.utg000033l.211.1
Phymatoceros_bulbiculosus_v1_contig17_g189240.t2
Phymatoceros_bulbiculosus_v1_contig22_g207630.t1
Phymatoceros_bulbiculosus_v1_contig28_g224480.t1
Anthoceros_angustus_002912
Anthoceros_agrestis_BONN_evm.model.Sc2ySwM_368.2609.1
Anthoceros_angustus_002909
Anthoceros_agrestis_BONN_evm.model.Sc2ySwM_368.2607.1
Anthoceros_agrestis_OXF_evm.model.utg000033l.204.1
Anthoceros_punctatus_evm.model.utg000041l.206.1
Anthoceros_angustus_002911
Leiosporoceros_dussii_v1_contig23_g60840.t1
Leiosporoceros_dussii_v1_contig23_g60860.t1
Leiosporoceros_dussii_v1_contig23_g60850.t1
Leiosporoceros_dussii_v1_contig23_g60810.t1
Leiosporoceros_dussii_v1_contig23_g60820.t1
Leiosporoceros_dussii_v1_contig23_g60790.t1
Leiosporoceros_dussii_v1_contig23_g60800.t1
Phaeoceros_carolinianus_v1_contig1_g39620.t1
Phaeoceros_sp_v1_contig4_g70330.t1
Phaeoceros_sp_v1_contig1_g00950.t1
Phaeoceros_carolinianus_v1_contig4_g98970.t1
Paraphymatoceros_pearsonii_v1_contig5_g24330.t1
Notothylas_orbicularis_v1_contig4_g174270.t1
Phymatoceros_bulbiculosus_v1_contig21_g205610.t1
Anthoceros_agrestis_BONN_evm.model.Sc2ySwM_368.1160.1
Anthoceros_agrestis_OXF_evm.model.utg000010l.214.1
Anthoceros_punctatus_evm.model.utg000151l.329.1
Anthoceros_angustus_010178
Anthoceros_fusiformis_v1_contig16_g182560.t1
Phymatoceros_bulbiculosus_v1_contig15_g181270.t1
Phaeoceros_carolinianus_v1_contig3_g77880.t1
Phaeoceros_sp_v1_contig7_g102820.t1
Phaeoceros_carolinianus_v1_contig11_g152410.t1
Paraphymatoceros_pearsonii_v1_contig55_g206470.t1
Notothylas_orbicularis_v1_contig1_g21870.t1
Phymatoceros_bulbiculosus_v1_contig15_g181250.t1
Phaeoceros_carolinianus_v1_contig3_g77890.t1
Phaeoceros_sp_v1_contig7_g102830.t1
Paraphymatoceros_pearsonii_v1_contig55_g206460.t1
Leiosporoceros_dussii_v1_contig8_g28670.t1
Phymatoceros_bulbiculosus_v1_contig15_g181280.t1
Phymatoceros_bulbiculosus_v1_contig15_g181290.t1
P_patens_Pp3c7_9420V3.1.p
M_polymorpha_Mapoly0012s0063.1.p
M_polymorpha_Mapoly0012s0071.1.p
M_polymorpha_Mapoly0012s0070.1.p
M_polymorpha_Mapoly0012s0068.1.p
M_polymorpha_Mapoly0012s0067.1.p
M_polymorpha_Mapoly0012s0062.1.p
M_polymorpha_Mapoly0012s0065.1.p
M_polymorpha_Mapoly0204s0001.1.p
M_polymorpha_Mapoly0932s0001.1.p
M_polymorpha_Mapoly0204s0002.1.p
M_polymorpha_Mapoly0204s0003.1.p
M_polymorpha_Mapoly0359s0002.1.p
M_polymorpha_Mapoly0008s0080.1.p
M_polymorpha_Mapoly0008s0097.1.p
M_polymorpha_Mapoly0008s0109.1.p
M_polymorpha_Mapoly0063s0074.1.p
M_polymorpha_Mapoly0079s0066.1.p
Bryophyte
α-CA
C_subellipsoidea_58562
C_subellipsoidea_67822
C_braunii_tr|A0A388KF32|A0A388KF32_CHABU
C_braunii_tr|A0A388KF39|A0A388KF39_CHABU
C_braunii_tr|A0A388MBQ9|A0A388MBQ9_CHABU
C_braunii_tr|A0A388KF52|A0A388KF52_CHABU
C_braunii_tr|A0A388LIZ3|A0A388LIZ3_CHABU
C_braunii_tr|A0A388KR04|A0A388KR04_CHABU
C_braunii_tr|A0A388KR39|A0A388KR39_CHABU
C_braunii_tr|A0A388KF63|A0A388KF63_CHABU
C_braunii_tr|A0A388KF41|A0A388KF41_CHABU
D_salina_Dusal.0018s00001.1.p
D_salina_Dusal.0852s00001.1.p
D_salina_Dusal.0049s00039.1.p
D_salina_Dusal.0049s00038.1.p
M_endlicherianum_ME000018S02371
M_endlicherianum_ME000018S02377
M_endlicherianum_ME000018S02372
V_carteri_Vocar.0025s0008.1.p
V_carteri_Vocar.0025s0010.1.p
V_carteri_Vocar.0025s0011.1.p
C_reinhardtii_Cre04.g223050.t1.2
C_reinhardtii_Cre04.g223100.t1.2
C_subellipsoidea_48539
K_nitens_kfl00242_0130_v1.1
Algal
α-CA
Anthoceros_angustus_008185
Anthoceros_fusiformis_v1_contig1_g15290.t1
Anthoceros_agrestis_BONN_evm.model.Sc2ySwM_344.4293.1
Anthoceros_agrestis_OXF_evm.model.utg000038l.124.1
Anthoceros_punctatus_evm.model.utg000124l.216.1
Phaeoceros_carolinianus_v1_contig6_g117600.t1
Phaeoceros_sp_v1_contig17_g177660.t1
Paraphymatoceros_pearsonii_v1_contig29_g146970.t1
Notothylas_orbicularis_v1_contig3_g111400.t2
Phymatoceros_bulbiculosus_v1_contig2_g42270.t2
Leiosporoceros_dussii_v1_contig8_g28130.t1
P_patens_Pp3c18_6840V3.1.p
S_fallax_Sphfalx08G147100.1.p
M_polymorpha_Mapoly0041s0087.1.p
Bryophyte
CAH3
K_nitens_kfl00210_0020_v1.1
C_reinhardtii_Cre09.g415700.t1.2
V_carteri_Vocar.0011s0335.1.p
D_salina_Dusal.0407s00003.1.p
M_endlicherianum_ME000241S03756
K_nitens_kfl00135_0080_v1.1
K_nitens_kfl00030_0090_v1.1
K_nitens_kfl00158_0230_v1.1
Algal
CAH3
0.4 substitutions/site

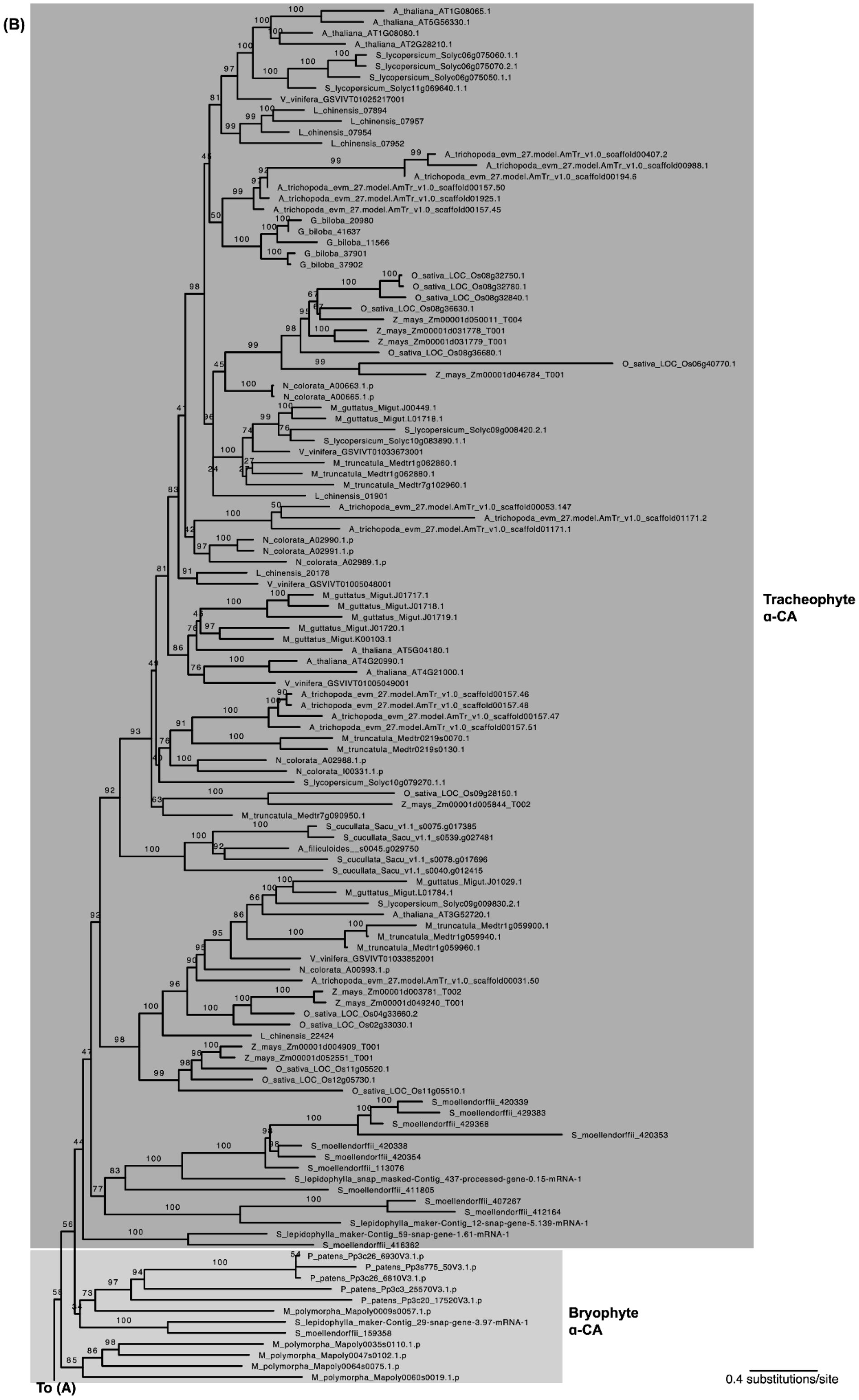


**Supplementary Fig. 5. Expanded CAH3 phylogeny.** Numbers above branches are bootstrap support values.

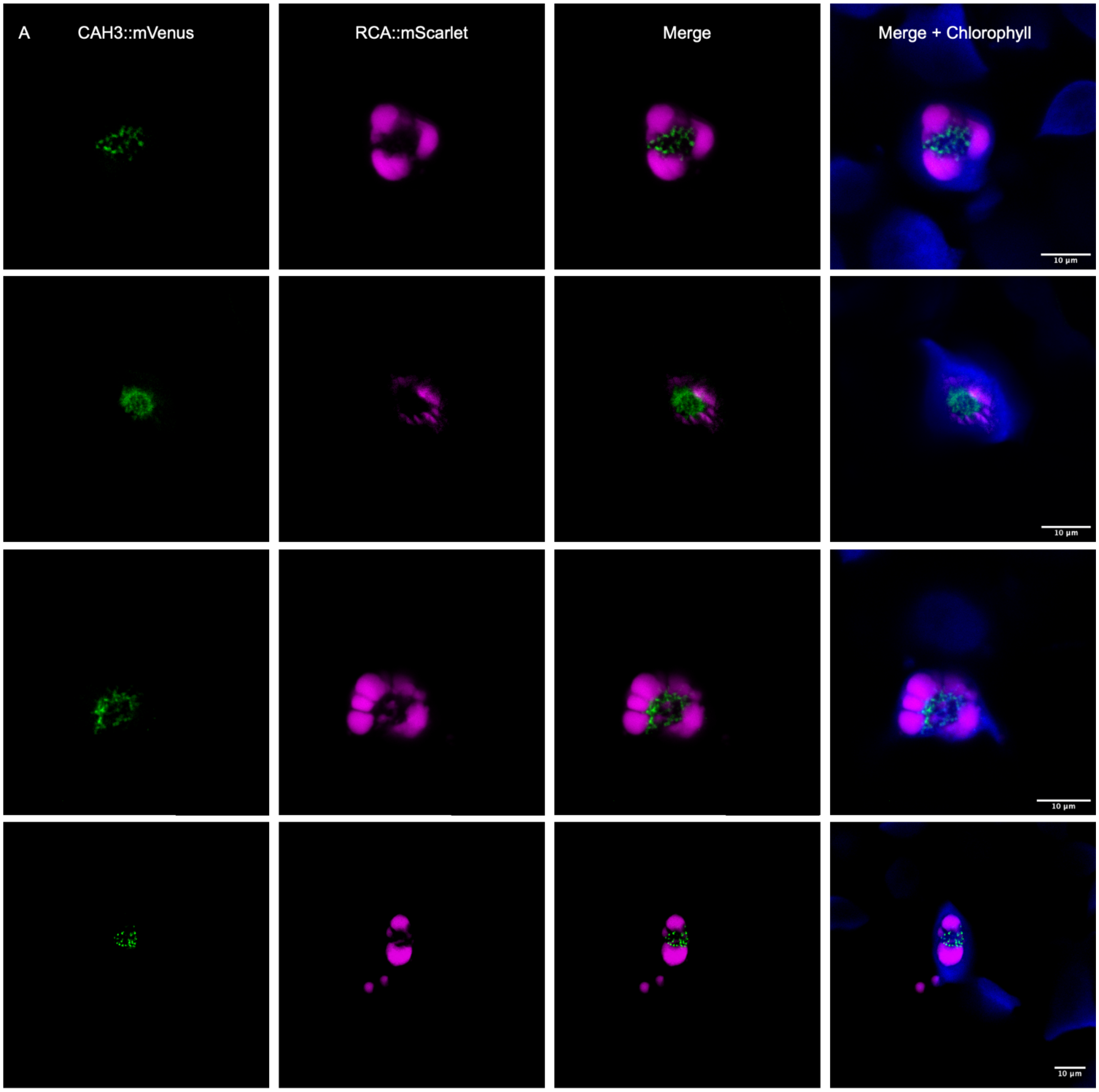


B

MGSLRVLVPSTLLGSNGQAESWRADLGTARPALPRRGGKGNAGGVRCCVREAGPADREEGE
GSKSSLIARDGLGRRGMLSAAMLATMTCVCTEGAEAAVARHAAVEQRRREALQLQTLQGAKA
RGGSRIKKLGGKAAVGTAEWEYGNSCGPGEWGSVCAAGRVQSPVNFEMNKVQEKGEKAMS
DLMFDYGPVQPTFLNTGHGTMQVNFPGGRNKLKIGDRVLDLLQF**H**F**H**TPSEHSFNGVHHTME
A**H**LVHGDPKTKSLAVVGVLLDAKSRVSANKALQAALEYSPKEHYKTARGPDNFSLSPSLLLPYA
GRIRKKRGYMYYQGSLTTPPCSEGVEWYVMETPVSISYAQVVEFMLYVGDSRTLALNTRPVQP
LGQRRVYRGPMQA

**Supplementary Fig. 6.** ***A. agrestis*** **CAH3.** (A) Additional images of CAH3::mVenus. CAH3::mVenus are shown in green, RCA::mScarlet in magenta, and chlorophyll autofluorescence in blue. (scale bar, 10 µm.) (B) Protein sequence of CAH3, lumenal transit peptide is underlined and conserved catalytic sites bolded.

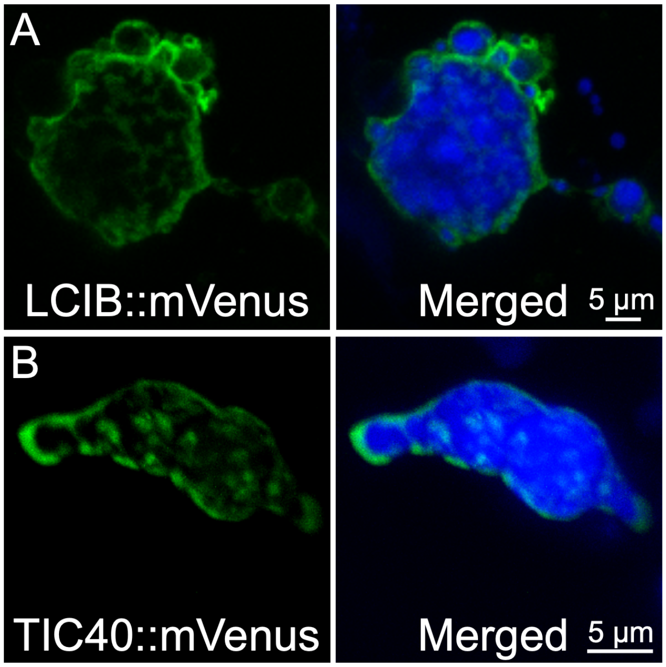


**Supplementary Fig. 7. *A. agrestis* LCIB and TIC40**. Images of *A. agrestis* separately expressing (A) LCIB::mVenus and (B) TIC40::mVenus. mVenus tagged proteins are shown in green, RCA::mScarlet in magenta and chlorophyll autofluorescence in blue. (Scale bar, 5 µm.)

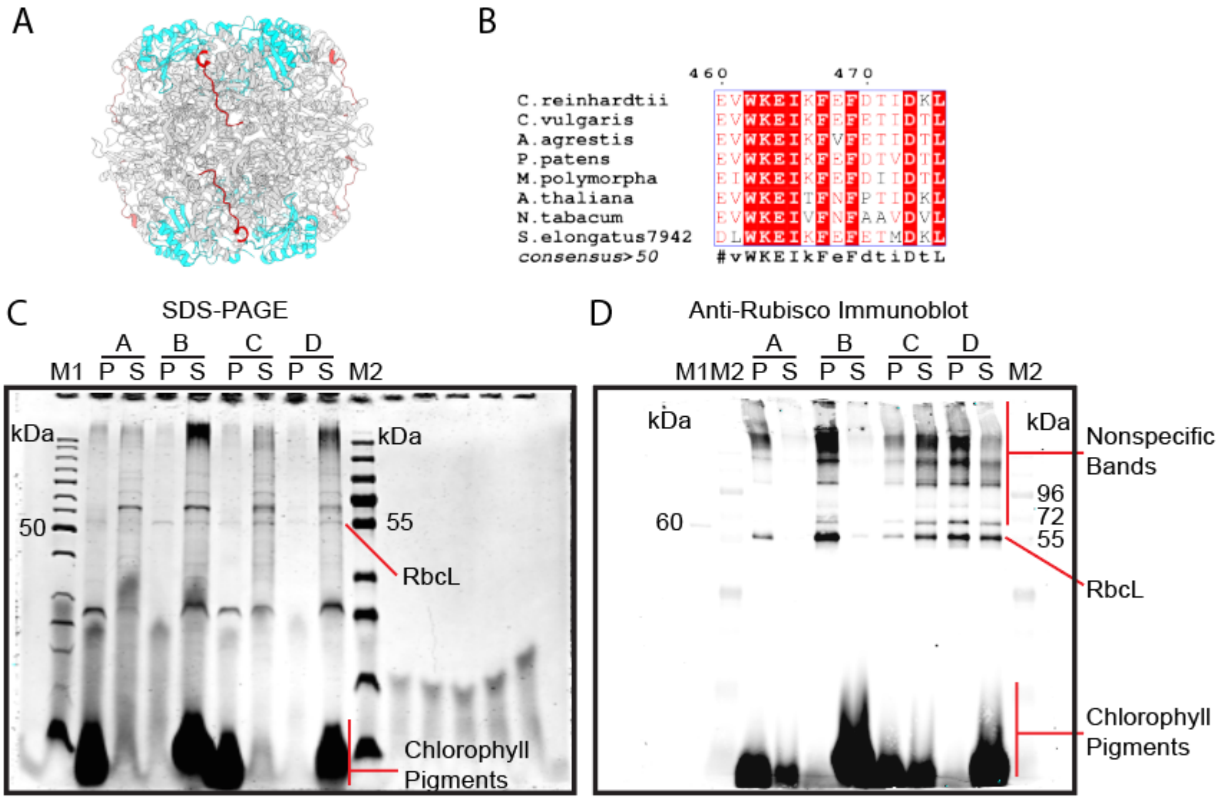


Buffer composition
A: 20 mM HEPES pH 7.0, 50 mM NaCl, 12.5% v/v glycerol
B: Buffer A + 2% v/v Triton-X 100
C: 20 mM CAPS pH 11.0, 50 mM NaCl, 12.5% v/v glycerol
D: Buffer C + 2% v/v Triton-X 100

pH of buffer after lysis
A - 5.5, B - 6, C - 8, D - 7
Soluble protein C/D>B>A
Insoluble protein B>D>A>C

**Supplementary Fig. 8. Rubisco antibody design and *A. agrestis* tissue lysis.** (A) Antibodies were raised against the *A. agrestis* Rubisco large subunit C-terminus sequence, highlighted in red in the Rubisco molecule (PDB: 2V63)[1]. Rubisco large and small subunits are shown in grey and blue respectively. (B) Alignment of Rubisco C-terminus sequences from *C. reinhardtii*, *Chlorella vulgaris*, *A. agrestis*, *Physcomitrium patens*, *Marchantia polymorpha*, *Arabidopsis thaliana*, *Nicotiana tabacum*, and *Synechococcus elongatus* PCC 7942. Alignment graphics were generated with ESPript[2]. (C) Lysis of *A. agrestis* tissue with different buffers. Insoluble, P, and soluble proteins, S, were analyzed using SDS-PAGE using 8–16% Mini-PROTEAN® TGX™ Precast Protein Gels. (D) Immunoblot was analyzed with antibodies raised against Rubisco C-terminus.

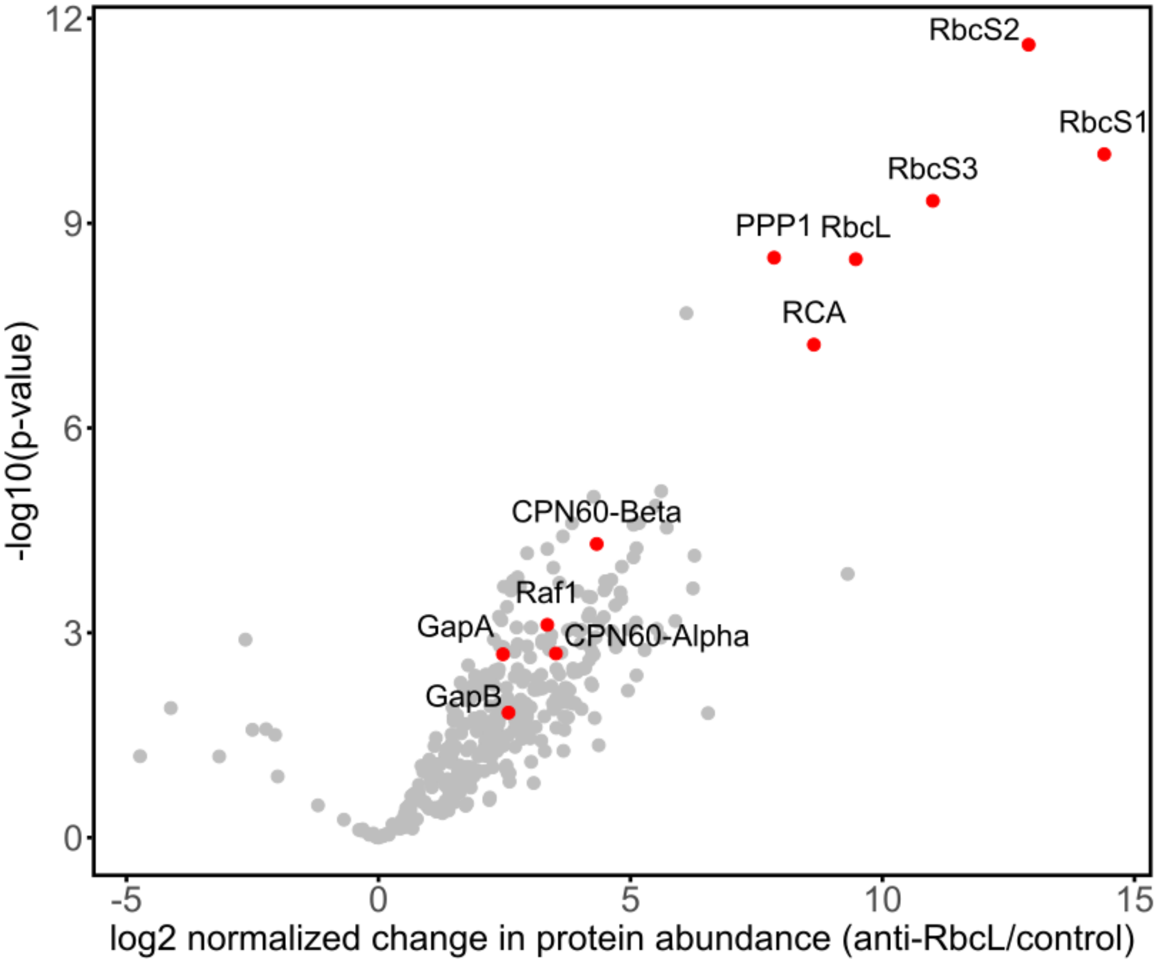


**Supplementary Fig. 9. Volcano plot of proteins identified in Rubisco co-IP.** Statistical significance of protein enrichment (-log10(p-value)) was plotted against the change in protein abundance of Rubisco co-IP samples compared to the control. Proteins of interest were colored in red with associated annotations displayed. A total of five technical replicates were each performed for Rubisco co-IP samples and the control.

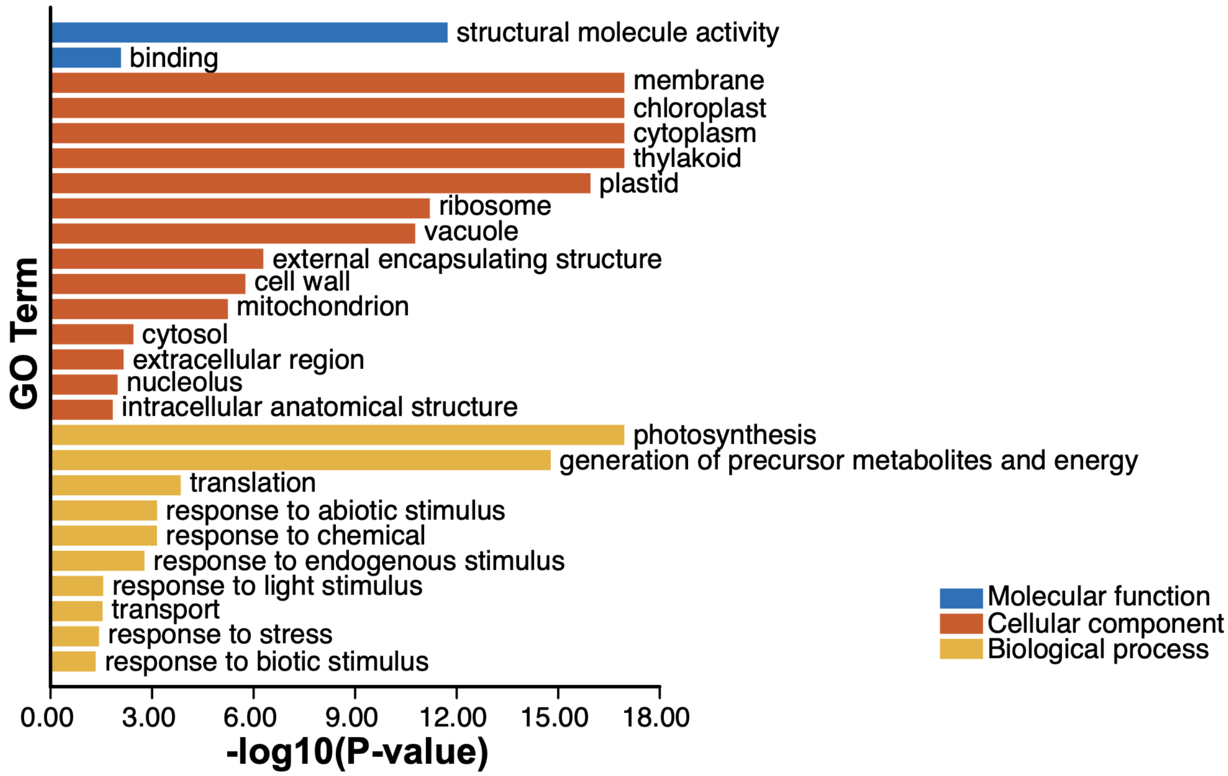


**Supplementary Fig. 10. Enriched gene ontology (GO) terms of proteins that co-immunoprecipitated with Rubisco.**

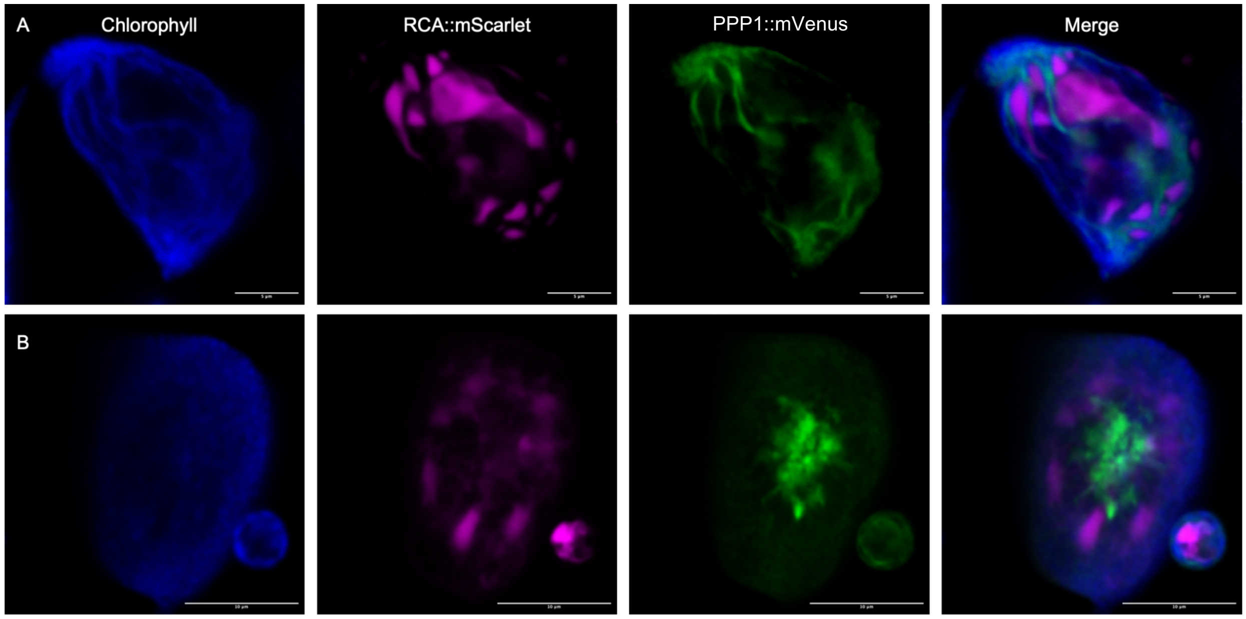


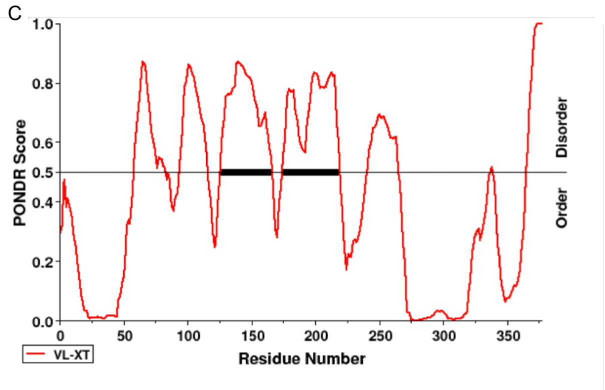


D

MAAVGCGPRLALSRRCGLQHQGVVHGRFKVAMAPAVGSTW
VWHAFPRISWVQEAAVRSKRDASAYASNGKENGPLDENGVE
KQGTWQDGKEGFDVESEAEKVLRIDEESQALLKDLAVAAAR
VSYLQSDLEKHKKREQEIANLRAEIERYAEEQAAADSAVAKY
RDEVADAEQELYEASIALVSARVGDSSEDISGKWELVTVDED
AERIESGKAATVSSLVGSVASLPFLLSERTDLPNLLPSLLSER
NVDLTENVDLSGVAGVAGLSGIAGVAGVFLSCALFGVCYRYIV
RRDLGNFQLKAGATAAFALVRGIAQVDTTQALVGFPDQGLSE
VLQACLDAGESLVTIAFAAVALDYCLREGLLEPFPSARSRRDK

**Supplementary Fig. 11. PPP1::mVenus localization.** PPP1::mVenus showed two localizations either to the (A) stroma or (B) thylakoid knots that are surrounded by pyrenoids. (scale bar, 5 and 10 µm). PPP1::mVenus are shown in green, RCA::mScarlet in magenta and chlorophyll autofluorescence in blue. (C) Disorder prediction (Pondr) of PPP1. (D) Protein sequence of PPP1, SMC domain is highlighted in blue.

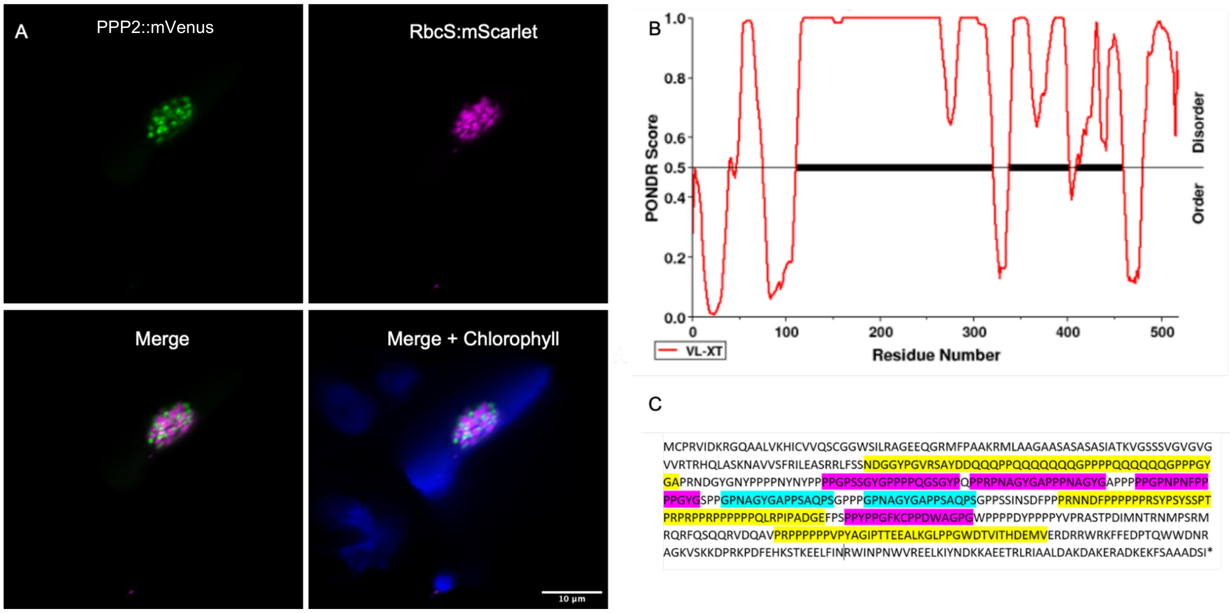


**Supplementary Fig. 12. PPP2::mVenus localization.** (A) PPP2::mVenus localized to the space between pyrenoid subunits. PPP2::mVenus are shown in green, RbcS::mScarlet in magenta and chlorophyll autofluorescence in blue. (scale bar, 10 µm.) (B) Disorder prediction (Pondr) of PPP2. (C) Protein sequence of PPP2, three repeat motifs are highlighted in yellow, magenta and teal respectively.

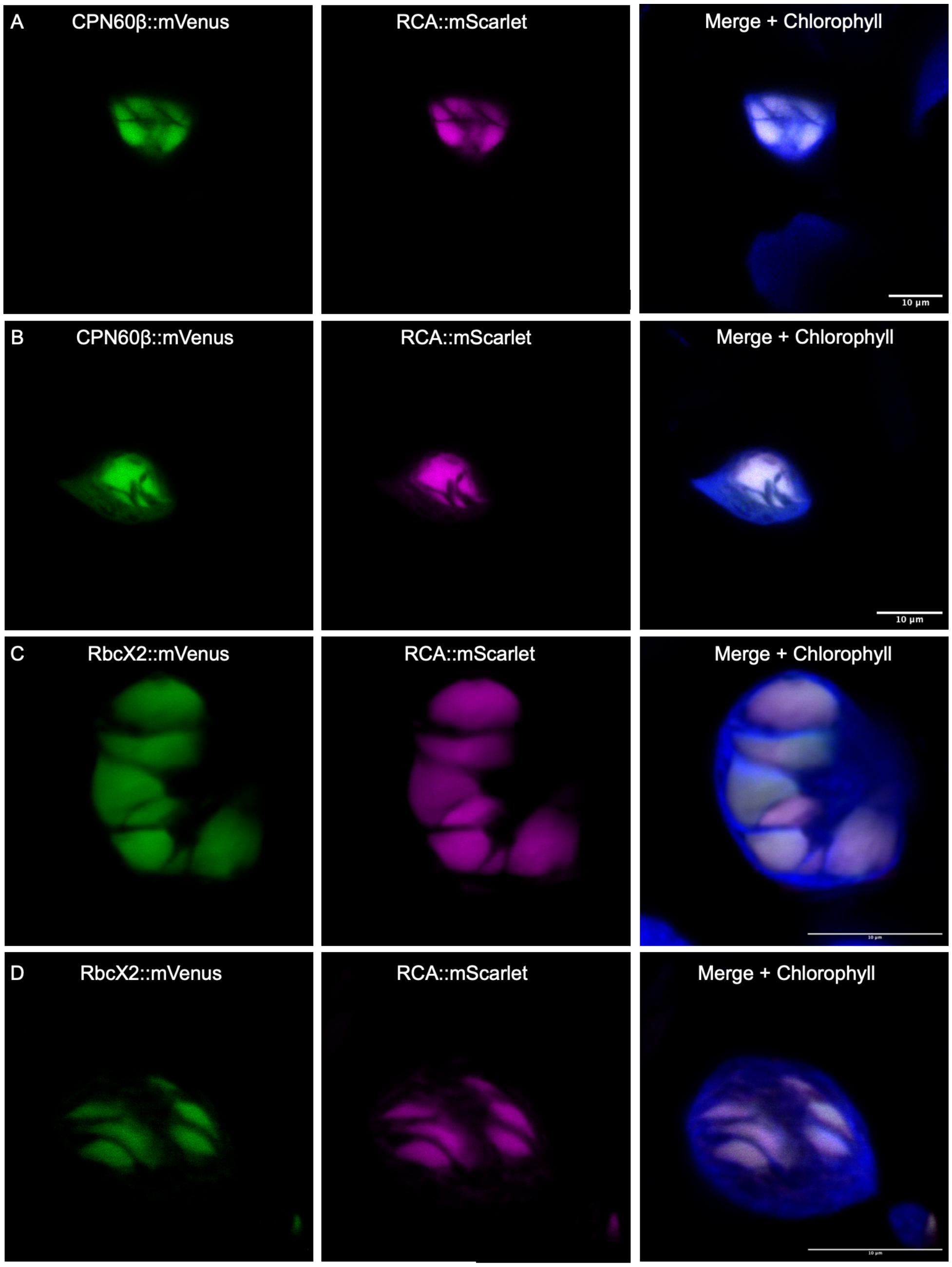


**Supplementary Fig 13.** ***A. agrestis*** **Cpn60β and RbcX2.** Additional images of *A. agrestis* expressing (A and B) Cpn60β::mVenus and (C and D) RbcX2::mVenus. mVenus tagged proteins are shown in green, RCA::mScarlet in magenta and chlorophyll autofluorescence in blue. (Scale bar, 10 μm.).

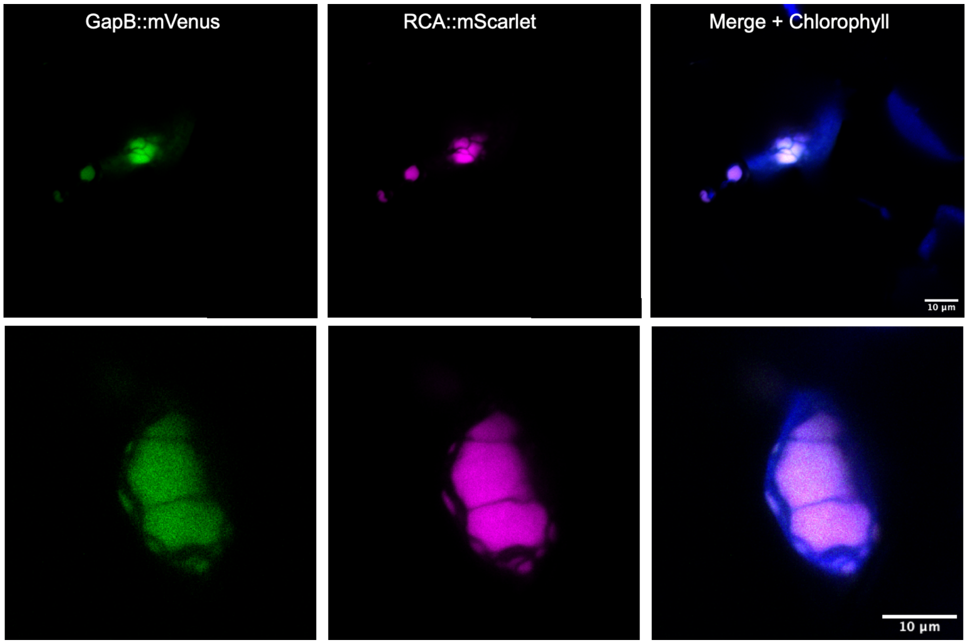


**Supplementary Fig 14. *A. agrestis* GapB.** Additional images of *A. agrestis* expressing GapB::mVenus. mVenus tagged proteins are shown in green, RCA::mScarlet in magenta and chlorophyll autofluorescence in blue. (scale bar, 10 µm.).

**Supplementary Table 1. List of gene IDs used in this study.**

| Gene | *Anthoceros agrestis* gene ID, version [3] | *A. agrestis* gene ID, version [4] | Chlamydomonas gene ID | Expression level in *A. agrestis* (transcripts per million) |
|---|---|---|---|---|
| RCA | AnagrOXF.S1G164600.t1 | AagrOXF_evm.model.utg000011l.50.1 | Cre04.g229300 | 709 |
| PSAH | AnagrOXF.S1G410400.t1 | AagrOXF_evm.model.utg000058l.57.1 | Cre07.g330250 | 2449 |
| PSBP | AnagrOXF.S1G127000.t1 | AagrOXF_evm.model.utg000063l.559.1 | Cre12.g550850 | 1739 |
| BST1 | AnagrOXF.S2G372500.t1 | AagrOXF_evm_utg000116l.141 | Cre16.g662600 (BST1)<br>Cre16.g663450 (BST2)<br>Cre16.g663400 (BST3) | 861 |
| BST2 | AnagrOXF.S1G176100.t1 | AagrOXF_evm.model.utg000080l.43.1 | Cre16.g662600 (BST1)<br>Cre16.g663450 (BST2)<br>Cre16.g663400 (BST3) | 8 |
| BST3 | AnagrOXF.S1G344800.t1 | AagrOXF_evm.model.utg000100l.403.1 | Cre16.g662600 (BST1)<br>Cre16.g663450 (BST2)<br>Cre16.g663400 (BST3) | 14 |
| BST4 | AnagrOXF.S2G189400.t1 | AagrOXF_evm.model.utg000042l.477.1 | Cre16.g662600 (BST1)<br>Cre16.g663450 (BST2)<br>Cre16.g663400 (BST3) | 5 |
| β-CA1 | AnagrOXF.S4G438500.t1 | AagrOXF_evm.model.utg000063l.200 | N.A. | 1519 |
| CAH3 | AnagrOXF.S2G141200.t1 | AagrOXF_evm.model.utg000038l.124.1 | Cre09.g415700 | 212 |
| LCIB | AnagrOXF.S3G266500.t3 | AagrOXF_evm.model.utg000033l.200.2 | Cre10.g452800 | 130 |
| PPP1 | AnagrOXF.S1G403800.t1 | AagrOXF_evm.model.utg000119l.172.1 | N.A. | 119 |
| PPP2 | AnagrOXF.S4G428300.t2 | AagrOXF_evm.model.utg000063l.301 | N.A. | 335 |
| Cpn60β | AnagrOXF.S4G153700.t2 | AagrOXF_evm.model.utg000091l.123.1 | Cre07.g339150 | 11 |
| RbcX2 | AnagrOXF.S1G289300.t1 | AagrOXF_evm.model.utg000049l.48.1 | Cre01.g030350 | 82 |
| GapB | AnagrOXF.S4G265100.t1 | AagrOXF_evm.model.utg000002l.202.1 | Cre01.g010900 | 446 |

**Supplementary Table 2. List of genomes used in Orthofinder.**

| | | |
|---|---|---|
| Vascular plants | Seed Plants | *Arabidopsis thaliana, Amborella trichopoda, Ginkgo biloba, Liriodendron chinense, Mimulus guttatus, Medicago truncatula, Nymphaea colorata, Oryza sativa, Solanum lycopersicum, Vitis vinifera, Zea Mays* |
| | Ferns | *Azolla filliculoides, Salvinia cucullata* |
| | Lycophyte | *Selaginella lepidophylla, Selaginella moellendorffii* |
| Bryophytes | Hornworts | *Anthoceros agrestis* (Oxford), *Anthoceros agrestis* (Bonn), *Anthoceros angustus, Anthoceros fusiformis, Anthoceros punctatus, Leiosporoceros dussii, Notothylas orbicularis, Paraphymatoceros pearsonii, Phaeoceros carolinianus, Phaeoceros spp, Phymatoceros bulbiculosus* |
| | Moss | *Fontinalis antipyretica, Pleurozium schreberi, Physcometrium patens, Sphagnum fallax* |
| | Liverworts | *Marchantia polymorpha* |
| Charophyte | | *Chara braunii, Klebsormidium patens, Mesotaenium endlicherianum, Penium margaritaceum* |
| Chlorophyte | | *Chlamydomonas reinhardtii, Coccomyxa subellipsoidea, Dunaliella salina, Micromonas pusilla, Ostreococcus lucimarinus, Volvox carteri* |
